# Mycobacterium tuberculosis LprE enhances bacterial persistence by inhibiting cathelicidin and autophagy in macrophages

**DOI:** 10.1101/230698

**Authors:** Avinash Padhi, Ella Bhagyaraj, Mehak Zahoor Khan, Mainak Biswas, Srabasti Sengupta, Geetanjali Ganguli, Manaswini Jagadev, Ananyaashree Behera, Kaliprasad Pattanaik, Ishani Das, Pawan Gupta, Vinay Kumar Nandicoori, Avinash Sonawane

**Author notes:** Corresponding author (AS).

## Abstract

*Mycobacterium tuberculosis(Mtb)* lipoproteins are known to facilitate bacterial survival by manipulating the host immune responses. Here, we have characterized a novel *Mtb* lipoprotein LprE(LprE_Mtb_), and demonstrated its role in mycobacterial survival. LprE_Mtb_ acts by down-regulating the expression of cathelicidin, Cyp27B1, VDR and p38-MAPK via TLR-2 signaling pathway. Deletion of *lprE_Mtb_* resulted in induction of cathelicidin and decreased survival in the host. Interestingly, LprE_Mtb_ was also found to inhibit autophagy mechanism to dampen host immune response. Episomal expression of LprE_Mtb_ in non-pathogenic *Mycobacterium smegmatis*(*Msm*) increased bacillary persistence by down-regulating the expression of cathelicidin and autophagy, while deletion of LprE_Mtb_ orthologue in *Msm*, had no effect on cathelicidin and autophagy expression. Moreover, LprE_Mtb_ blocked phago-lysosome fusion by suppressing the expression of EEA1, Rab7 and LAMP-1 endosomal markers by down-regulating IL-12 and IL-22 cytokines. Our results indicate that LprE_Mtb_ plays an important role in mycobacterial pathogenesis in the context of innate immunity.

## INTRODUCTION

Tuberculosis (TB), caused by *Mycobacterium tuberculosis (Mtb)*, is a widespread disease that kills more than 1.5 million people every year worldwide [1]. Macrophages, the primary resident cells of *Mtb*, are important components of innate defense mechanisms of the host. Macrophages play a crucial role in recognition, phagocytosis, and killing of invaders. Non-pathogenic mycobacteria such as *M. smegmatis* (*Msm*) are readily killed by the macrophages, whereas pathogenic mycobacteria (*Mtb*) employ various strategies to survive inside the macrophages like prevention of phago-lysosome fusion [2], autophagy inhibition [3], modulation of host cytokine production [4], inhibition of reactive oxygen and nitrogen species [5] and the manipulation of antigen presentation to prevent or alter the quality of T-cell responses [6].

Among the key effector molecules responsible for bacterial killing are antimicrobial peptides such as cathelicidin and defensins that are expressed in different cells such as neutrophils, macrophages, monocytes and epithelial cells [7]. In contrast to the multiple defensins, only one cathelicidin gene, *CAMP* (cathelicidin antimicrobial peptide), has been found in humans [8, 9]. The gene product human cationic antimicrobial peptide-18 (hCAP18) is transcribed from the *CAMP* gene that contains vitamin D response elements (VDRE) in its promoter and requires enzymatic digestion by human neutrophil proteinase 3 (PR3) to produce mature LL-37 that exhibit broad antimicrobial activity [10]. LL-37 is an amphipathic α-helical peptide that binds to negatively charged groups of the bacterial outer membrane causing disruption of the bacterial cell wall [11]. Previously, we and other studies have shown that LL-37 is able to kill both non-pathogenic and pathogenic mycobacteria both *in vitro* and *in vivo* conditions [12, 13, 14].

Previously, it has been shown that vitamin D strongly up regulates the expression of cathelicidin [15]. The proposed model suggested that toll-like receptor-2/1 (TLR2/1) activation of macrophages induce the expression of CYP27B1 (25-hydroxyvitamin D-1α-hydroxylase) which, in turn, leads to the production of bioactive 1, 25(OH)_2_D from circulating inactive 25(OH) D [16]. The production of 1, 25(OH)_2_D activates *CAMP* gene expression through vitamin D receptor [17].

Bacterial lipoproteins are known to perform diverse functions such as host cell adhesion, cell invasion, nutrient transport, drug resistance and evasion of host defense mechanisms [18]. For example, *Streptococcus pneumoniae* PsaA lipoprotein is involved in the colonization of host cells and antibiotic resistance [19,20]. *Borrelia burgdorfori* VlsE lipoprotein is involved in bacterial persistence in host cells [21] and *Haemophilus influenzae* P6 lipoprotein activate host immune cells by inducing the secretion of pro-inflammatory cytokines through TLR-2 binding [22]. Similarly, bacterial lipoproteins have been shown to stimulate proliferation of B cells leading to increased immunoglobulin secretion [23]. Thus, bacterial lipoproteins are able to activate both the innate and adaptive wings of the immune system. Mycobacterium genome encodes for 99 lipoproteins, however, the functions of many of them are still unknown. Several of them have been shown to play a major role in virulence [16, 24, and 25]. *Mtb* PstS-1, a 38 kDa lipoprotein, is involved in phosphate transport as well as in inducing apoptosis [25]. Deletion mutants of *Mtb lgt* and *lsp* showed a significant reduction in virulence [26]. Similarly, an extensively studied *Mtb* 19-kDa lipoprotein (LpqH) has been shown to strongly induce the innate immunity by activating cathelicidin mediated autophagy in a TLR-2 dependent manner [24,16]. LpqH binds to TLR-2 that leads to activation of cytokines such as interleukin-12 (IL-12), interleukin-1 beta (IL-1β) and tumor necrosis factor-alpha (TNF-α). Moreover, LpqH also plays a pivotal role in bacterial survival by inhibiting interferon-gamma (IFN-γ) response genes such as *CIITA* (Class II trans activator) that leads to reduced antigen presentation [24]. Another study has shown that a synthetic 19-kDa *Mtb* derived lipopeptide enhances the antimicrobial capacity of monocytes via TLR2/1 signaling, vitamin D and VDR-dependent pathway [27]. This involved induction of the *CAMP* gene and its protein [28]. Some of the lesser studied mycobacterial lipoprotein genes include LprG, LprA, LpqB, and LpqM.

In the present study, we have shown that one of the *Mtb* lipoproteins, LprE (LprE_Mtb_), is involved in intracellular bacterial survival and evasion of host immune responses. We demonstrate that LprEMtb enhances bacillary survival by inhibiting the expression of *CAMP* via TLR-2-p38-Cyp27B1-VDR signaling pathway in human macrophages. We also demonstrate that LprE_Mtb_ downregulates the pathogen induced IL-1β pro-inflammatory response. Furthermore, mechanistic studies showed that LprE_Mtb_ inhibits autophagy and phago-lysosome fusion by down-regulating the expression of several endosomal markers such as Early Endosome Antigen 1 (EEA1), Rab7 and lysosomal-associated membrane protein 1 (LAMP-1), and IL-12 and IL-22 cytokines, thus aid bacterial survival in human macrophages. In summary, we have characterized a novel *Mtb* lipoprotein that aids bacterial survival in host macrophages by down-regulating the expression of antimicrobial peptide cathelicidin.

## RESULTS

### Genetic organization of LprE in *M. tuberculosis* genome

Initial annotation from the Tuberculist Web server suggested that *Mtb* H37Rv LprE (LprE_Mtb_), encoded by a 609 bp *Rv1252c* gene, belongs to a previously uncharacterized Rv_dir301 operon (webTB.org database). This operon consists of yet another uncharacterized *Rv1251c* gene (**Fig 1A**). In *Mtb* CDC1551, a clinical isolate, LprE is encoded by an *MT1291* gene (**Fig 1B**). Multiple sequence alignment and BLAST results showed 100% sequence homology between *Mtb* H37Rv and *Mtb* CDC1551 *LprE* genes (**Suppl Fig 1**).The tertiary structure of LprE is not available. Therefore, we predicted its structure using a Modeller program. The LprE sequence showed high structural homology (94.5%) with human mitochondrial RNA polymerase (PDB ID: 3SPA) (**Fig 1C**). Domain analysis by ProDom program (http://www.http://prodom.prabi.fr/) showed the presence of a conserved lipoprotein binding domain at amino acids 81-201 position (data not shown). To further confirm that LprE is a lipoprotein, we aligned LprE_Mtb_ with *Escherichia coli* outer membrane lipopolysaccharide transport lipoprotein LptE (PDB ID: 4NHR). We observed 100% homology between lipoprotein binding domains of LprE_Mtb_ and *E. coli* LptE proteins (**Fig 1D**).

**Fig 1:**
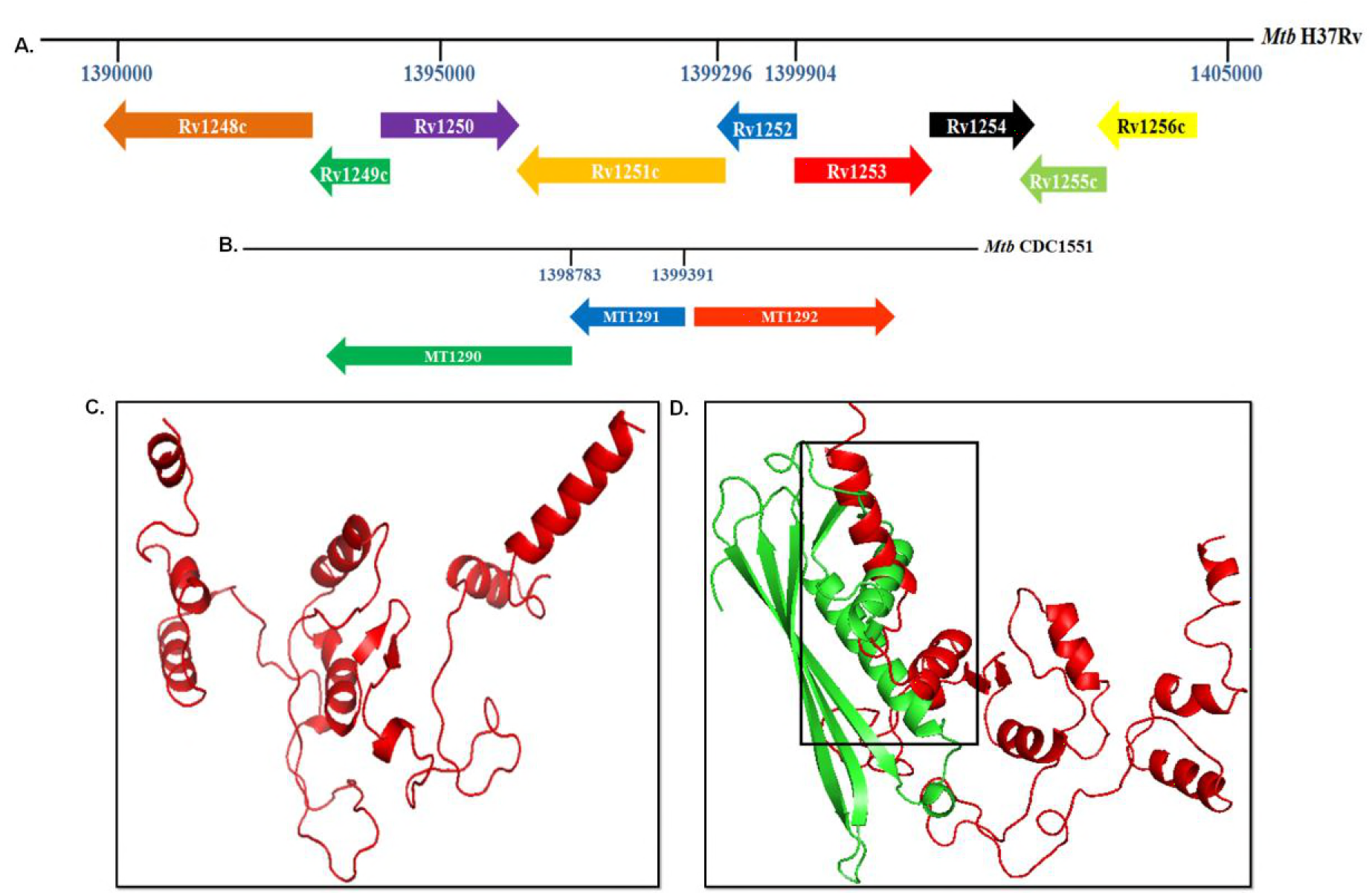
*In silico* analysis of LprE_Mtb_. **A**. Genetic organization of *Rv1252c* in *Mtb* H37Rv genome. **B**. Genetic organization of *MT1291* in *Mtb* CDC1551 genome. **C**. Prediction of LprE_M_t_b_ tertiary structure with modeler v9.1.2 using Human Mitochondrial RNA Polymerase (PDB ID: 3SPA) as a template. **D**. Domain alignment of energy minimized structure of LprE_Mtb_ and *E. coli* outer membrane lipopolysaccharide transport lipoprotein LptE (PDB ID: 4NHR).

### Deletion of *LprE* in *M. tuberculosis* CDC1551 decreased bacterial survival in human macrophages

Pathogenic mycobacteria employ various strategies to survive and replicate inside the host cells. Earlier reports showed that *Mtb* lipoproteins play a crucial role in the intracellular survival of bacteria and that deletion of lipoproteins decrease *Mtb* virulence [26, 29]. Therefore, we investigated the role of LprE_Mtb_ in bacterial survival. Towards this, THP-1 cells were infected with *Mtb*, *Mtb*Δ*LprE*, and the complemented *Mtb*Δ*LprE::LprE* strains. Cells were lysed at 12, 24, 48 and 72 h post infection and the intracellular bacterial survival was determined. While *Mtb* and *Mtb*Δ*LprE::LprE* strains showed a time dependent increase in intracellular bacterial burden, significant decrease was observed when THP1 cells were infected with *Mtb*Δ*LprE* (**Fig 2A**). These results suggest a pivotal role for LprE_Mtb_ in the survival of *Mtb* inside macrophages.

**Fig 2:**
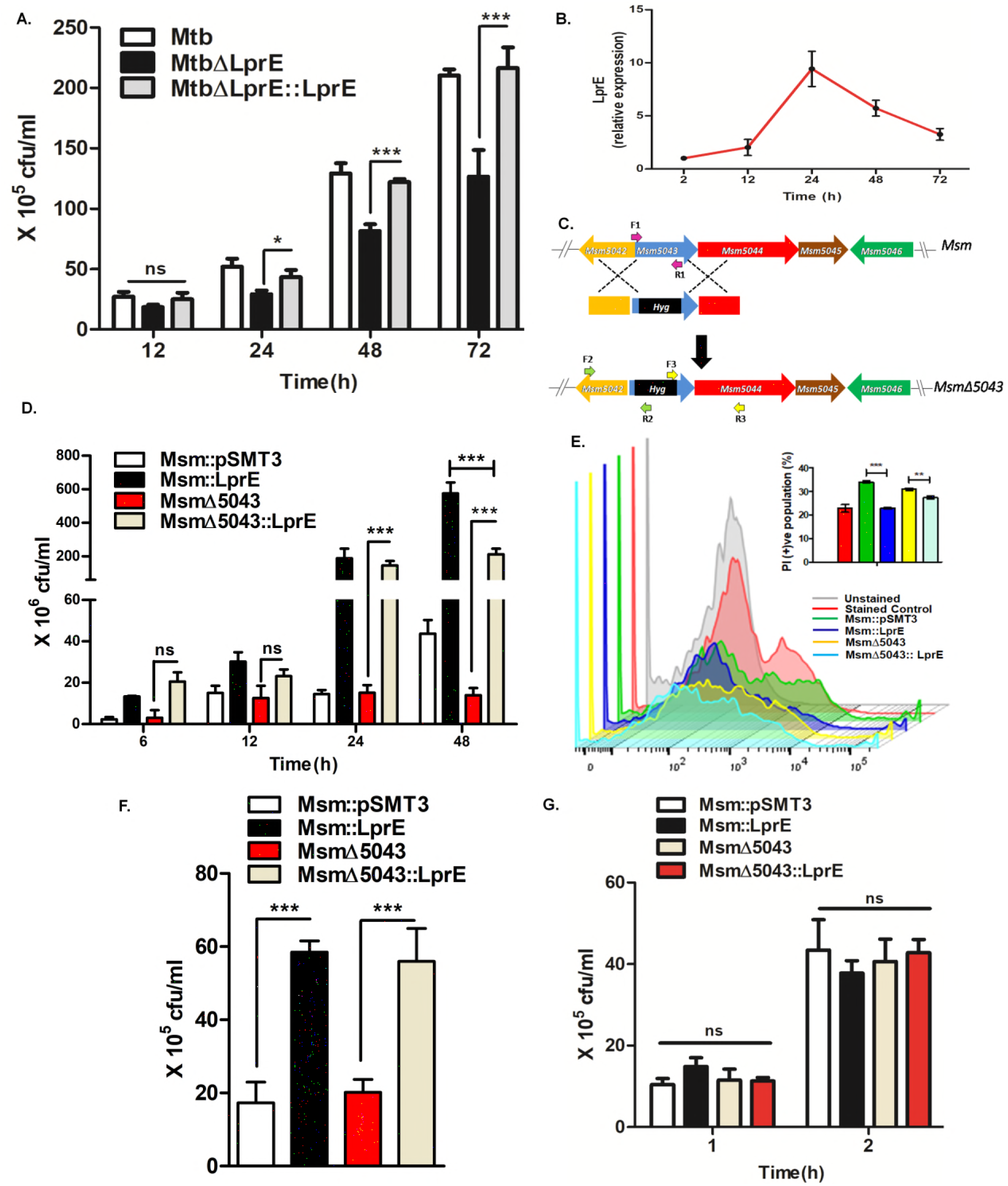
Role of LprE_Mtb_ in intracellular survival in human macrophages. **A**. THP-1 cells were infected with *Mtb, Mtb*Δ*LprE* and *Mtb*Δ*LprE::LprE* complemented strain. The cells were lysed and the intracellular survival was determined at 12, 24, 48 and 72 h post infection by cfu assay. B. qRT-PCR analysis of *LprE_Mtb_* expression was performed from intracellular bacteria isolated from macrophages after 2, 12, 24, 48 and 72 h post infection. The expression values were normalized with *Msm sigA* gene. **C**. Schematic representation of construction of *Msm*Δ*5043* mutant (*MSM_5043* in *Msm*). The primers used for screening knockout are represented by arrows. **D**. THP-1 cells were infected with *Msm::pSMT3, Msm::LprE, Msm*Δ*5043* and *MsmΔ5043::LprE* strains. The cells were lysed and the intracellular survival was determined at 6, 12, 24, and 48 h post infection by cfu assay. **E**. THP-1 cells were infected with recombinant *Msm* strains for 2 h and intracellular survival was determined at 24 h post infection. Intracellular bacteria were stained with propidium iodide (PI) and analyzed by Flow cytometer. PI positive population was considered as dead bacteria, while negative as live. **F**. Human PBMCs were isolated from healthy individuals, differentiated into macrophages with GM-CSF and infected with *Msm::pSMT3, Msm::LprE, MsmΔ5043* and *MsmΔ5043::LprE* strains for 2 h. The intracellular count was determined 24 h post infection by cfu assay. **G**. Phagocytosis rates of the above mentioned *Msm* strains was determined by determining intracellular counts in THP-1 cells at early time points of 1 and 2 h post infection. Experiments were performed in triplicates; Mean ± SD; *** for P<0.0001, ** for P<0.001, * for P< 0.05; ns, non-significant.

To further confirm the role of LprE in bacterial survival, LprE_Mtb_ was episomally expressed in non-pathogenic *Msm* (*Msm::LprE*) with the help of pSMT3 shuttle vector. Several studies including our previous work have used *Msm* as a surrogate model to study the role of various *Mtb* proteins in pathogenesis [30–34]. The qRT-PCR analysis confirmed that *LprE* is ectopically expressed in *Msm* strain grown under *in-vitro* condition (**Fig 2B**). As shown *lprE_Mtb_* level reached a maximum at 24 h followed by a gradual decrease in expression level upto 72 h. The transcript levels were normalized with *Msm* housekeeping *sigA* at 2 h and the fold changes were calculated.

*Msm* genome analysis showed presence of *MSMEG_5043,* an ortholog of LprE_Mtb_ [35]. To preclude the effect of *MSMEG_5043*, we first constructed *Msm*Δ*5043* mutant by allelic exchange method (**Fig 2C**). Deletion of *MSMEG_5043* was confirmed by PCR using gene specific and flanking region primers (**Suppl Fig 2A**). *Msm*Δ*5043* strain was electroported with pSMT3-LprE_Mtb_ construct to generate a *Msm*Δ*5043::LprE* complementation strain. We did not observe significant difference in the *in vitro* growth pattern of *Msm::pSMT3, Msm::LprE, MsmA.5043* and *Msm*Δ*5043::LprE* strains (**Suppl Fig 2B**). These results are consistant with the growth pattern observed for *Mtb, Mtb*Δ*LprE*, and *Mtb*Δ*LprE::LprE* strains (data not shown), suggesting that the presence or absence *LprE_Mtb_* does not impact on the bacterial growth kinetics. In contrast, when compared with *Msm::pSMT3* and *Msm*Δ*5043*, the intracellular survival of *Msm::LprE* and *Msm*Δ*5043::LprE* strains in THP1 cells was 10-15 fold higher after 24 and 48 h of postinfection (**Fig 2D**). However, intracellular count of *Msm*Δ*5043::LprE* was observed to be 3 fold less when compared to *Msm::LprE* 48 h post infection (p<0.0001; **Fig 2D**). To corroborate above results, we determined the intracellular survival of bacteria 24 h post infection with the help of flow cytometry. Cells were harvested 24 h post infection, lysed to release the intracellular bacteria and stained with propidium iodide (PI), a non-permeable nucleic acid binding dye that binds specifically to dead or with disintegrated cell wall bacteria. Flow cytometry analysis showed less PI stained *Msm::LprE* and *Msm*Δ*5043:: LprE* bacteria as compared with *Msm::pSMT3* and *Msm*Δ*5043* strains (**Fig 2E**), indicating less death in LprE_Mtb_ expressing *Msm* strains.

Consistent with above results *Msm::LprE* and *Msm*Δ*5043::LprE* also showed increased intracellular survival in human monocyte derived macrophages as compared to *Msm::pSMT3* and *Msm*Δ*5043* strains (**Fig 2F**). We did not observe any difference in the phagocytosis rate of *Msm::pSMT3, Msm::LprE, Msm*Δ*5043* and *Msm*Δ*5043::LprE* strains (**Fig 2G**) indicating that the increased survival of *Msm::LprE* and *Msm*Δ*5043::LprE* strains is not due to the difference in the uptake of bacteria by macrophages.

### LprE_Mtb_ inhibits cathelicidin expression in human macrophages

Macrophages exhibit antimicrobial response by inducing the expression of cationic antimicrobial peptide cathelicidin (hCAP18/LL-37) [36]. However, *Mtb* is known to down regulate the expression of cathelicidin to avoid killing by macrophages [37]. Previously, it has been shown that 19-kDa lipoprotein from *Mtb* interacts with TLR-2, which subsequently up-regulates the expression of Cyp27B1 hydroxylase, VDR translocation into the nucleus and finally the induction of cathelicidin expression [14]. Our previous study also showed that cathelicidin is able to kill both pathogenic and non-pathogenic mycobacteria under *in vitro* and *ex vivo* conditions [13]. Therefore, we first evaluated the expression of *CAMP*, which encodes human cathelicidin, in *Mtb, Mtb*Δ*LprE*, and *Mtb*Δ*LprE::LprE* infected THP-1 cells. We observed an ~4-fold increase in *CAMP* expression in cells infected with *MtbΔLprE* compared with *Mtb* and *Mtb*Δ*LprE::LprE* infected cells (**Fig 3A**). In contrast, *Msm::LprE* and *Msm*Δ*5043::LprE* infection of THP1 cells resulted significant down-regulation of *CAMP* expression when compared with *Msm::pSMT3* and *Msm*Δ*5043* infected cells (**Fig 3B**). These results suggest that LprE_Mtb_ is involved in the inhibition of cathelicidin expression, which in turn promotes the bacterial survival. To further validate the role of cathelicidin, we evaluated bacterial survival in THP-1 cells treated with 50 μg/ml of LL-37 2 h before (pre-treatment) and after (post-treatment) the infection. Earlier we have shown that treatment with cathelicidin (50 μg/ml) would eliminate *Mtb* under *in vitro* and *ex vivo* conditions without exhibiting any toxic effect on macrophages [13]. Furthermore we established that *Msm* elimination is dependent on cathelicidin expression [13]. In agreement with our previous data, we observed increased killing of *Msm::LprE* and *Msm*Δ*5043::LprE* under both pre and post-treated conditions in comparison with untreated cells (**Fig 3C**). However, more bacterial killing was observed under post-treated condition. As expected treatment with 50 μg/ml LL-37 did not show any cytotoxic effect on THP-1 cells (data not shown).

**Fig 3:**
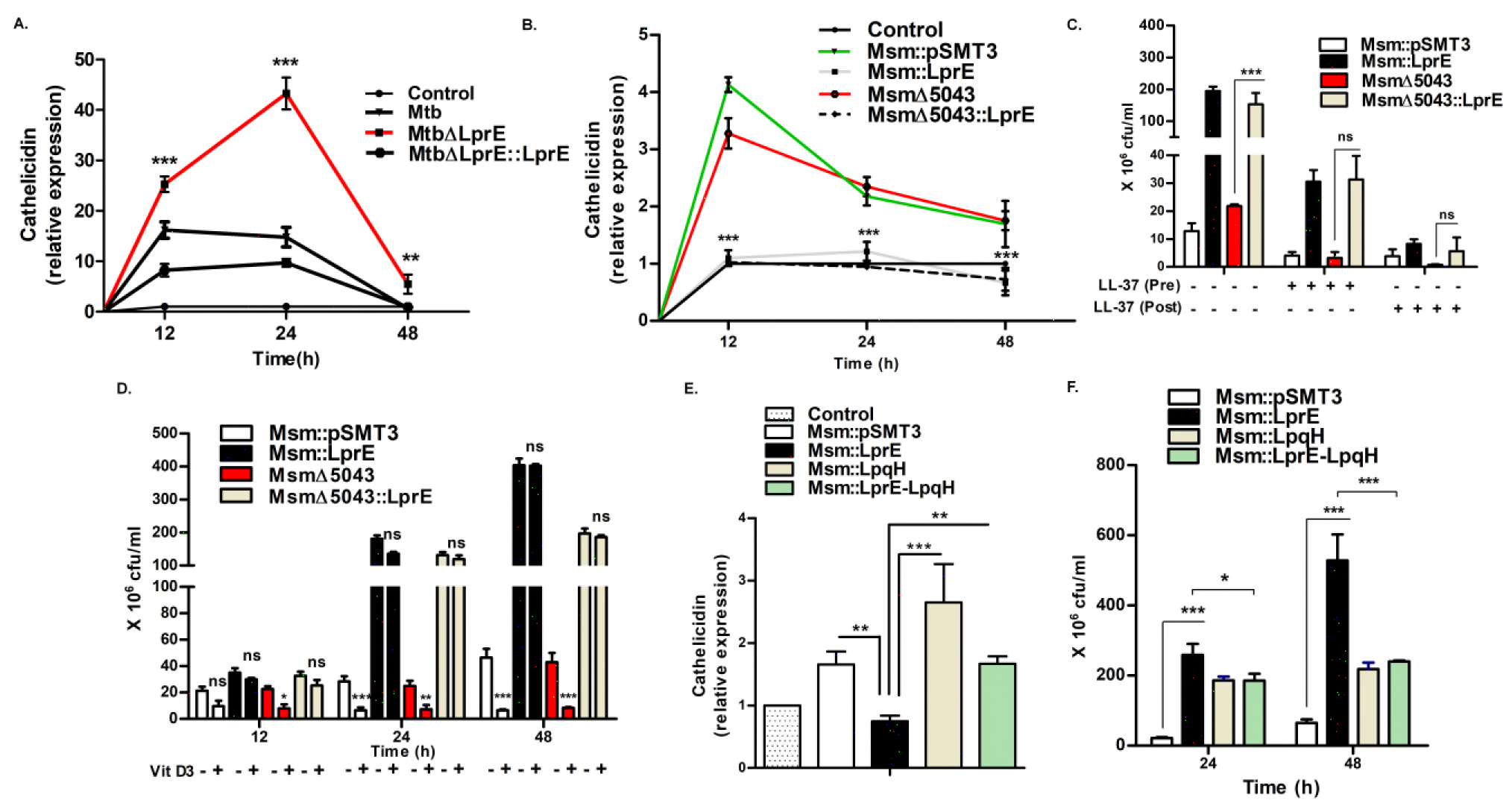
LprE_Mtb_ inhibits the expression of cathelicidin. **A**. THP-1 cells were infected with *Mtb, Mtb*Δ*LprE* and *Mtb*Δ*LprE::LprE* strains. The cells were harvested at 12, 24 and 48 h post infection, RNA was isolated and converted to cDNA. Cathelicidin (*CAMP*) expression was determined by qRT-PCR. **B.** Cathelicidin expression was determined by qRT-PCR in THP-1 cells infected with *Msm::pSMT3, Msm::LprE, Msm*Δ*5043* and *Msm*Δ*5043:LprE* at indicated time points. **C.** Cells were treated with 50 μg/ml LL-37 2 h before (Pre) and 2 h post (Post) infection. Intracellular counts of *Msm::pSMT3, Msm::LprE, Msm*Δ*5043* and *Msm*Δ*5043::LprE* were determined 24 h post infection in untreated, pre and post treated cells by cfu assay. **D.** Survival of *Msm::pSMT3, Msm::LprE, Msm*Δ*5043* and *Msm*Δ*5043::LprE* in 1,25 D3-treated and untreated THP-1 cells 24 h post infection. Cells were treated with 10^-7^ M 1,25 D3 18 h before infection. **E and F.** THP-1 cells were infected with *Msm::pSMT3, Msm::LprE, Msm::LpqH* and *Msm* co expressing *LprE_Mtb_* and *LpqH_Mtb_ (Msm::LprE-LpqH)*. 24 h post infection cells were harvested, **(E)** *CAMP* expression was checked by qRT-PCR and **(F)** intracellular survival was determined by cfu assay. Experiments were performed in triplicates; Mean ± SD; *** for P<0.0001, ** for P<0.001, * for P< 0.05; ns, non-significant.

**Fig 3:**
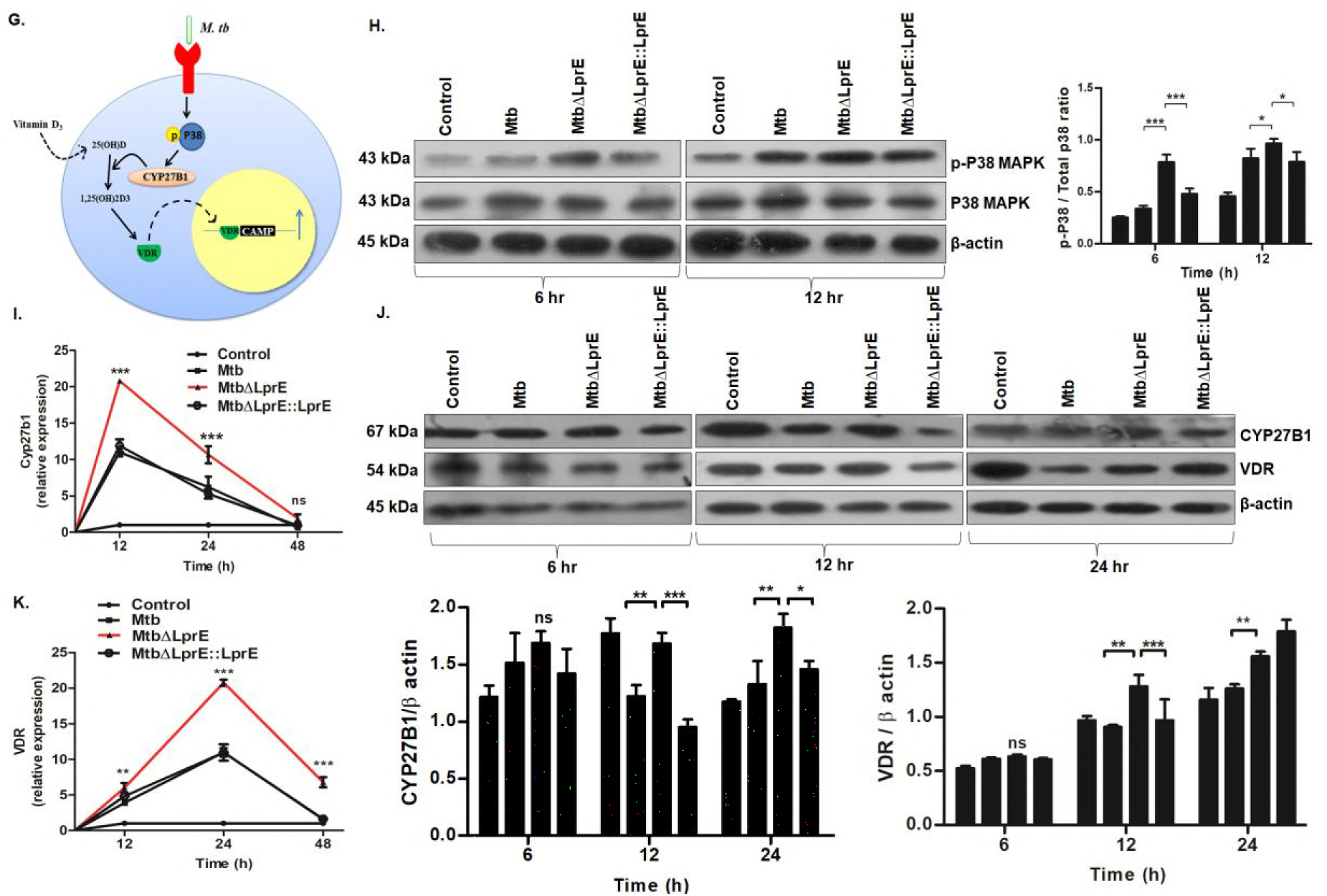
*Mtb*Δ*LprE* infection derepresses the expression of cathelicidin via p38 MAPK pathway in macrophages. **G**. Mycobacterial antigens activate p38 MAPK dependent Cyp27B1 and Vitamin D3 which then binds to VDR to activate the transcription of *CAMP* gene. THP-1 cells infected with *Mtb, Mtb*Δ*LprE* and *Mtb*Δ*LprE::LprE* strains were harvested at indicated time points to isolate total RNA and proteins. **H**. Western blot analysis of p38 MAPK expression was performed using anti-phospsho-p38 antibody. The expression was normalized to total p38 MAPK and β actin as loading control. Expression of Cyp27B1 at transcriptional (**I**) and translational (**J**) levels 12, 24 and 48 h post infection was determined by qRT-PCR and Western blotting, respectively. Transcriptional (**K**) and translational (**J**) levels of Vitamin D Receptor (VDR) was determined in infected THP-1 cells. Quantitative densitometry of phosphor-p38 expression was done using Image J software under baseline conditions. The expression values were normalized with *GAPDH* in case of qRT-PCR and β actin in western blotting. Experiments were performed in triplicates; Mean ± SD; *** for P< 0. 0001, ** for P < 0.001, * for P < 0.05; ns, non-significant.

Vitamin D3 is a known inducer of cathelicidin, which results in increased bacterial killing in human macrophages [38]. As shown in Figure 3D, treatment with 20 nM Vitamin D3 further reduced *Msm::pSMT3* and *Msm*Δ*5043* survival compared with untreated cells (**Fig 3D**), whereas we observed no difference in the cfu’s of *Msm::LprE* and *Msm*Δ*5043::LprE* (**Fig 3D**), suggesting that LprE_Mtb_ impedes the vitamin D3 mediated response.

### LpqH_Mtb_ and LprE_Mtb_ antagonostically regulate *CAMP* expression

Mycobacterium genome encodes for 90 lipoproteins. Among them, 19-KDa lipoprotein LpqHMtb has been shown to induce cathelicidin expression to restrict intracellular mycobacterial growth [16]. Therefore, we investigated the impact of LpqH_Mtb_ on the modulation of *CAMP* expression in presence of LprE_Mtb_. Towards this, we co-expressed LpqH_Mtb_ and LprE_Mtb_ in *Msm* to generate a recombinant *Msm::LprE-LpqH* strain. *Msm* expressing LpqH_Mtb_ (*Msm::LpqH*) and LprE_Mtb_ *(Msm::LprE)* were used as control strains. In agreement with the previous report, we also observed increased *CAMP* expression in *Msm::LpqH* infected cells [16], whilst *Msm::LprE* infection inhibited *CAMP* expression. However, infection with LpqH_Mtb_ and LprE_Mtb_ co-expressing *Msm* strain (*Msm::LprE-LpqH*) showed the *CAMP* level in between LpqH_Mtb_ and LprE_Mtb_ expressing *Msm* strains (**Fig 3E**), suggesting that both LpqH and LprE are strong regulators of cathelicidin expression. Moreover, *Msm::LpqH* and *Msm::LprE-LpqH* strains showed decreased survival in comparison with *Msm::LprE* strain after 24(p<0.05) and 48 (p<0.0001) h of infection (**Fig 3F**).

### LprE_Mtb_ inhibits cathelicidin via p38 MAPK pathway in macrophages

Based on above observations, our subsequent studies were focused on the investigation of the molecular mechanism(s) that are responsible for the down-regulation of *CAMP* expression by LprE_Mtb_. It has been reported that LpqH_Mtb_ and TLR-2 interaction regulate *CAMP* expression via activation of downstream p38 MAPK signaling pathway [14]. Mycobacterial antigen-TLR 2 interaction activates p38 MAPK signaling pathway dependent modulation of 25-hydroxyvitamin D–1α-hydroxylase (CYP27B1), which convert inactive 25(OH) D to active form 1,25(OH)_2_D of vitamin D [27]. The latter then binds to vitamin D receptor (VDR) to activate the transcription of the *CAMP* gene (**Fig 3G**). We observed a significant increase in the expression of phospho-p38 in *Mtb*Δ*LprE* infected cells when compared with *Mtb* and *Mtb*Δ*LprE::LprE* at 6 h post-infection, however, no such significant difference was observed at 12 h post-infection (**Fig 3H**) indicating that LprE_M_t_b_ modulates phospho-p38 during the early infection process. Similarly, phospho-p38 expression was found to be moderately decreased in *Msm::LprE* and *Msm*Δ*5043::LprE* infected macrophages as compared with *Msm*Δ*5043* at 6 h post infection (**Suppl Fig 3A**). This is in agreement with the previous study that *Mtb* infection down-regulates phospho-p38 [39].

### LprE_Mtb_ down-regulates CYP27B1 and VDR expression in macrophages

Next, we studied the expression of cathelicidin regulation pathway intermediates CYP27B1 and the VDR. Increased expression of CYP27B1 was observed at both transcriptional (**Fig 3I**) and translational levels (**Fig 3J**) in *Mtb*Δ*LprE* infected macrophages as compared to *Mtb* and *Mtb*Δ*LprE::LprE* infected cells. Of note, CYP27B1 expression was found to be increased both at 12 and 24 h post-infection (**Fig 3J**). On the other hand, the expression of CYP27B1 was found to be significantly down-regulated at both transcriptional (**Suppl Fig 3B**) and translational (**Suppl Fig 3C**) levels in *Msm::LprE* and *Msm*Δ*5043::LprE* infected cells relative to *Msm::pSMT3* and *Msm*Δ*5043* infected macrophages.

Similarly, the expression of VDR was found to be significantly up-regulated at both transcriptional (**Fig 3K**) and translational (**Fig 3J**) levels in *Mtb*Δ*LprE* infected macrophages as compared to *Mtb* and *Mtb*Δ*LprE::LprE* infected cells. In contrast, infection with *Msm::LprE* and *Msm*Δ*5043::LprE* strains down-regulated VDR expression at both transcriptional (**Suppl Fig 3D**) and translational (**Suppl Fig 3E**) levels. Altogether these results indicate that LprE_Mtb_ inhibits the expression of *CAMP* through down-regulation of p38-Cyp27B1-VDR signaling pathway to augment bacterial survival in macrophages.

### LprE_Mtb_ down-regulate the expression of *CAMP* via TLR-2

Previous studies have shown that lipoproteins regulate cathelicidin expression through TLRs [40,41]. To identify the involvement of specific TLR in the regulation of *CAMP* expression, we first determined *in-silico* binding efficiency of LprE_Mtb_ with human TLR-1,2,4 and 6. For this, we first performed docking analysis of energy minimized LprE_Mtb_ protein with human TLR’s tertiary structure obtained from PDB database. LprE_Mtb_ showed strong binding efficiency with TLR-2 (-582.21) as compared to TLR-1 (-223.5), TLR-4 (2.89) and TLR-6 (-10.59) (**Suppl Fig 4A**), as determined by atomic contact energy (ACE) score obtained from PatchDock analysis [42].

To validate the *in-silico* data, we silenced TLR’s using siRNAs specific to human TLR-1,2,4 and 6 in THP-1 cells. Semi-quantitative RT-PCR analysis showed more than 80% knock-down efficiency in siRNA treated cells (**Fig 4A**). Then untreated, scrambled and TLR specific siRNA treated THP-1 cells were infected with *Msm::pSMT3*, *Msm*Δ*5043* and *Msm*Δ*5043::LprE* bacteria and intracellular bacterial survival were determined. Interestingly, we observed a substantial decrease in the survival of *Msm*Δ*5043::LprE* in TLR-2 silenced macrophages as compared with TLR-1, 4 and 6 silenced macrophages (**Fig 4B**). Moreover, *Msm*Δ*5043::LprE* bacterial count was found to be approximately 1.5-fold less in TLR-1 siRNA treated cells in comparison with TLR-4 and 6 silenced macrophages (**Fig 4B**). This correlates well with the high ACE score observed in docking studies of LprE_Mtb_ with TLR-1 in comparison to TLR-4 and 6. We also checked the expression of *CAMP*, *Cyp27B1* and *VDR* in TLR-1,4,2 and 6 silenced plus *Msm::pSMT3*, *Msm*Δ*5043* and *Msm*Δ*5043::LprE* infected macrophages. We observed significant increase in the expression of *CAMP*, *Cyp27B1* and *VDR* (**Fig 4C-E**) in *Msm*Δ*5043::LprE* infected samples when TLR-2 was silenced in comparison with scrambled siRNA treated and untreated cells. No difference in the expression of *CAMP* was observed in TLR-1,4 and 6 silenced macrophages (**Suppl Fig 4B**). Consistent with the above data *Msm*Δ*5043::LprE* infection still showed down-regulation of *CAMP* in TLR-1, 4 and 6 silenced macrophages (**Suppl Fig 4B**). These results unambiguously suggest that LprE_Mtb_ regulates *CAMP* expression through TLR-2 signaling pathway.

**Fig 4:**
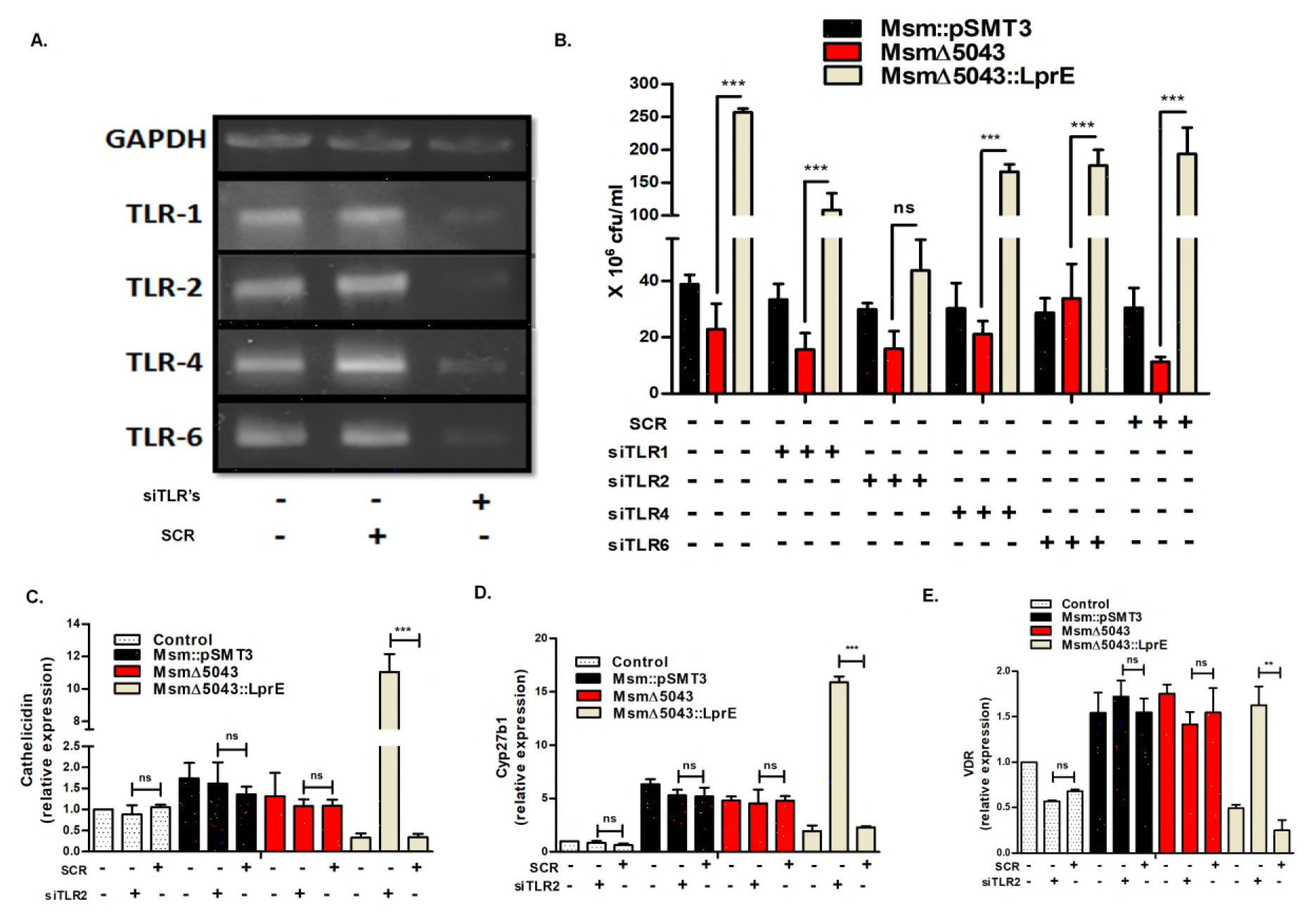
Expression of *CAMP*, *Cyp27b1, and VDR*, is regulated via TLR-2. **A**. Gene silencing was perfomed to knock down TLR-1,2,4 and 6. THP-1 cells were subjected to siRNA treatment by electroporation. Gene silencing was confirmed by RT-PCR (mRNA) and compared with cell treated with scrambled siRNA using gene specific primers. **B**. Untreated, TLR specific siRNA and scrambled siRNA treated THP-1 cells were infected with *Msm::pSMT3, Msm::LprE, Msm*Δ*5043* and *Msm*Δ*5043::LprE* strains. Cells were harvested at 24 h post infection and intracellular survival was determined by cfu assay. Untreated, siTLR-2 and scrambled treated THP-1 cells were infected with above mentioned strains. Cells were harvested at 24 h post infection, total RNA isolated, converted to cDNA and qRT-PCR was performed to determine the expression of **C**. Cathelicidin (CAMP), **D**. *Cyp27b1* and E. *VDR*. Experiments were performed in triplicates; Mean ± SD; *** for P < 0.0001, ** for P < 0.001; ns, non-significant.

### LprE_Mtb_ inhibits the production of pro-inflammatory cytokine IL-1β

IL-1β cytokine induces the expression of antimicrobial peptides through TLR signaling to clear the bacterial infection [43]. Active TB patients showed reduced levels of IL-1β suggesting that it may have a protective role in TB infection [44]. Next, we investigated the expression of IL-1β in *Mtb*, *Mtb*Δ*LprE*, and *Mtb*Δ*LprE::LprE* infected macrophages. qRT-PCR analysis showed significant up-regulation of IL-1β level in *Mtb*Δ*LprE* infected macrophages in comparison with *Mtb* and *Mtb*Δ*LprE::LprE* infected cells after 12 (p<0.0001) and 24 h (p<0.0001) of infection (**Fig 5A**). However, no difference in the expression levels of IL-1β was observed after 48 h of infection. In contrary, LprE_M_t_b_ expressing *Msm::LprE* and *Msm*Δ*5043::LprE* infected cells showed reduced expression of IL-1β at both 12 and 24 h post-infection (**Fig 5B**).

**Fig 5:**
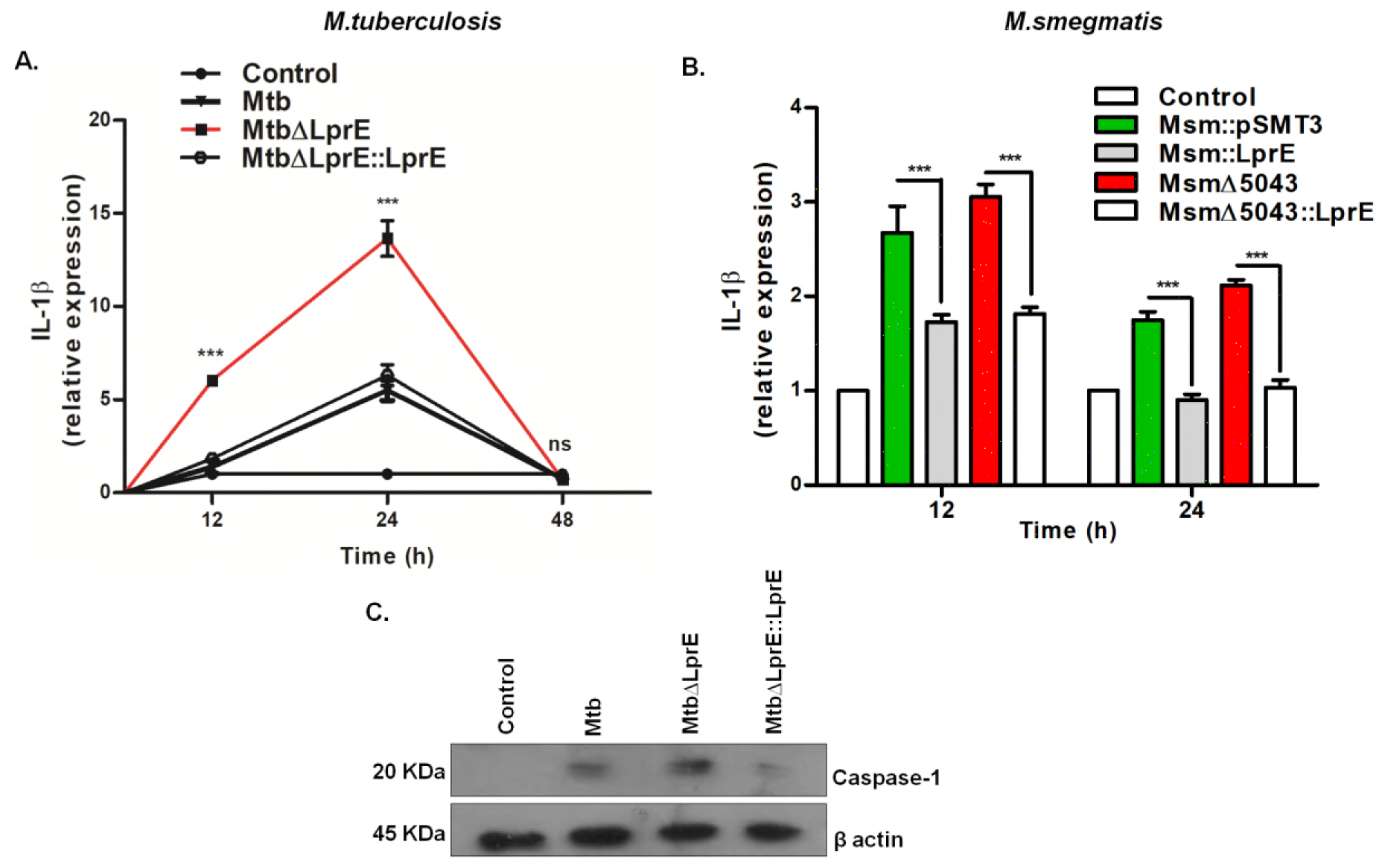
LprE_Mtb_ suppresses caspase-1 dependent IL-1β production. **A**. THP-1 cells infected with *Mtb, MtbΔLprE* and *MtbΔLprE::LprE* strains were harvested at the indicated time points to isolate total RNA. qRT-PCR using gene specific primers was performed to determine the expression of IL-1β. **B**. IL-1β expression was determined in THP-1 cells infected with *Msm::pSMT3, Msm::LprE, Msm*Δ*5043* and *Msm*Δ*5043::LprE* strains after 12 and 24 h of infection. **C**. Western blot to determine the expression of caspase-1 was done using an antibody against cleaved caspase-1 in THP-1 cells infected with *Mtb, Mtb*Δ*LprE* and *Mtb*Δ*LprE::LprE* strains 12 h post infection. Experiments were performed in triplicates. The data is shown from one representative experiment. Mean ± SD; *** for P < 0.0001; ns, nonsignificant.

In macrophages, processing and release of active IL-1β are dependent on caspase-1 activation [45]. Therefore, we investigated whether LprE_M_t_b_ mediated IL-1β down-regulation is dependent on caspase activation. Indeed, western blot analysis showed an increased level of cleaved caspase-1 in *Mtb*Δ*LprE* infected macrophages when compared with *Mtb* and *Mtb*Δ*LprE::LprE* infected cells after 12 h of infection (**Fig 5C**). Together data suggests that LprE_M_t_b_ suppresses caspase-1 dependent IL-1β production to facilitate bacterial persistence in macrophages.

### LprE_Mtb_ inhibits autophagy to dampen host immune response

Autophagy is a known host defense mechanism that plays an important role in the restriction of *Mtb* growth [46. To investigate if LprE_Mtb_ modulates autophagy, we evaluated the expression of several autophagic markers such as microtubule-associated protein 1 light chain 3 (LC3), ATG-5 and Beclin-1 in *Mtb, Mtb*Δ*LprE* and *Mtb*Δ*LprE::LprE* infected THP-1 cells. Western blot analysis showed the increased conversion of LC3-I to a characteristic autophagic induction marker LC3-II in *Mtb*Δ*LprE* infected cells (**Fig 6A**).We also examined the expression of an autophagic flux marker p62(SQSTM1) [47]. As shown in **Fig 6A**, p62 level decreased in case of *Mtb*Δ*LprE* infected macrophages (**Fig 6A**). Moreover, the expression of other autophagic markers such as Atg-5 and Beclin-1, which are recruited to the phagosomal compartments during autophagic vesicle formation [48], were also found to be increased in *Mtb*Δ*LprE* infected cells as compared to *Mtb* and *Mtb*Δ*LprE::LprE* infected cells (**Fig 6B**).

**Fig 6:**
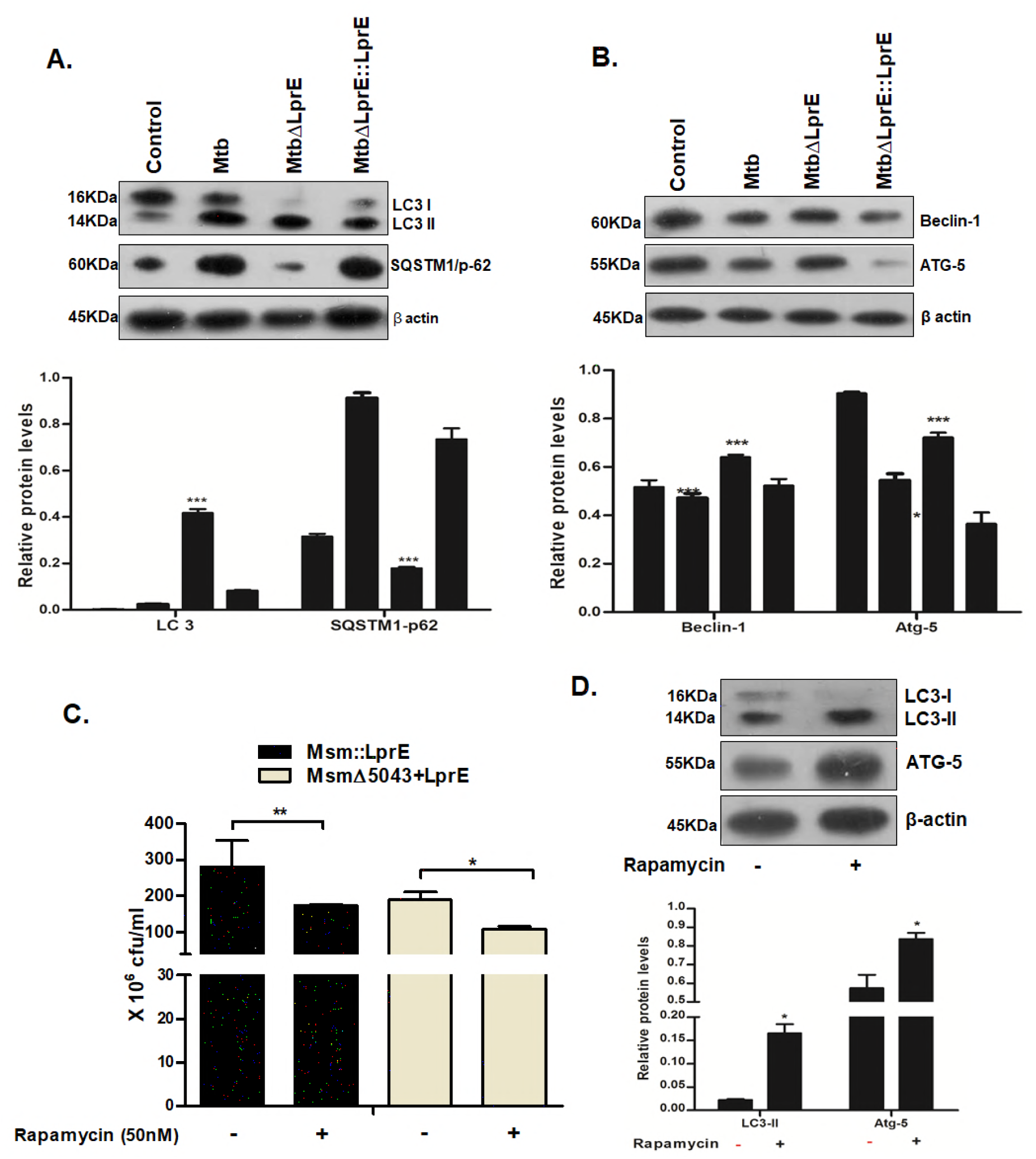
LprE_Mtb_ downregulates autophagy to mediate bacterial survival. THP-1 cells were infected with *Mtb, Mtb*Δ*LprE* and *MtbΔLprE::LprE* strains. Infected cells were harvested at respective time points and isolated proteins were analysed by western blotting. Western blot analysis of autophagy related proteins **A**. LC3 and SQSTM1/p62 at 6 h post infection and **B**. Atg-5 and Beclin-1 at 24 h was done using monoclonal antibodies against respective proteins. Densitometry values were calculated using Image J software to obtain a relative ratio of LC3II/LC3I. LC3II ratio, p62, Atg-5 and beclin-1 levels were normalized to β actin levels. **C.** Intracellular survival of *Msm::LprE* and *Msm*Δ*5043::LprE* 24h post infection in untreated and THP-1 cells pre-treated with rapamycin (50nM) as determined by cfu assay. Rapamycin treatment was confirmed by **D.** LC3II/LC3I ratio and Atg-5 expression. Experiments were performed in triplicates; Mean ± SD; *** for P < 0.0001, ** for P < 0.001, * for P < 0.05; ns, non-significant.

To further confirm the role of LprE_Mtb_ in autophagy inhibition, we also checked the expression of LC3, p62, Atg-5 and Beclin-1 in *Msm::pSMT3*, *Msm::LprE*, *Msm*Δ*5043* and *Msm*Δ*5043::LprE* infected macrophages. Consistent with above results, infection with both LprE_Mtb_ expressing *Msm::LprE* and *Msm*Δ*5043::LprE* strains significantly reduced the expression of LC3, Atg-5 and Beclin-5 (**Suppl Fig 5A & B**). On the other hand, p62 expression was found to be upregulated under similar conditions. Rapamycin is a known inducer of autophagy mechanism. Next, we checked if autophagy induction by rapamycin treatment exerts any impact on the outcome of bacterial survival. For this, THP-1 cells were pre-treated with rapamycin (50 nM) followed by infection with *Msm::LprE* and *Msm*Δ*5043::LprE* strains and the intracellular bacterial count was determined after 24 h of infection. Rapamycin treatment significantly reduced the survival of *Msm::LprE* and *Msm*Δ*5043::LprE,* which otherwise showed more survival by down-regulating the expression of autophagy, as compared to untreated cells (**Fig 6C**). We confirmed that rapamycin treatment indeed induced autophagy by determining the expression of LC3-II and Atg-5 by western blots analysis (**Fig 6D**).

### LprE_Mtb_ blocks phago-lysosome fusion by down-regulating the expression of endosomal markers

Non-pathogenic mycobacteria containing phagosomes readily fuse with lysosomes leading to the elimination of pathogen; however, pathogenic mycobacteria inhibit the fusion of phagosomes with lysosomes to survive in macrophages for an extended period of time. Previously, we have shown that exogenous administration of cathelicidin increased the colocalization of *M. bovis*-BCG containing phagosomes with lysosomes resulting in increased bacterial killing [11]. Based on these observations, we extended our study to investigate whether LprE_Mtb_ also affects the phagosome maturation and subsequently phago-lysosome fusion. For this, we checked the expression of early endosomal antigen 1 (EEA1), Rab7 and lysosomal associated marker protein-1 (LAMP-1) proteins, which are the hallmarks of early, late and lysosomal compartments, in THP-1 cells infected with *Mtb, Mtb*Δ*LprE,* and *Mtb*Δ*LprE::LprE* strains. It has been shown that recruitment of these marker proteins on endocytic compartments is necessary for the sequential maturation of endosomes and phagolysosome fusion [49,50]. Western blot analysis showed a significant increase in the levels of EEA1, Rab7 and LAMP-1 in *Mtb*Δ*LprE* infected cells in comparison to *Mtb* and *Mtb*Δ*LprE::LprE* infected cells (**Fig 7A &B**). On the other hand, macrophages infected with LprE_M_t_b_ expressing *Msm* strains showed decreased expression of these endosomal markers (**Suppl Fig 6A**).

**Fig 7:**
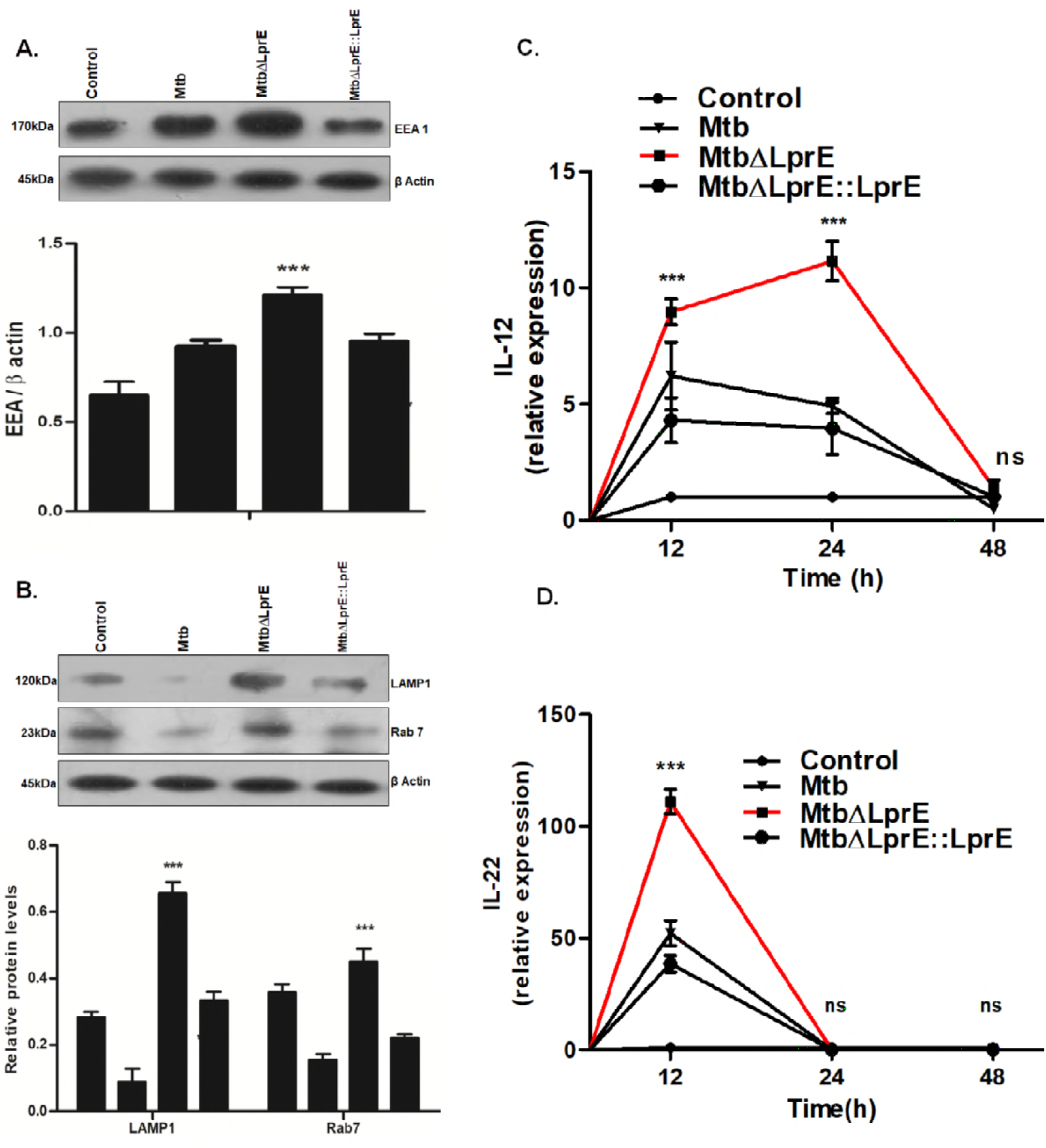
LprE_Mtb_ mediated phagosomal marker downregulation. THP-1 cells were infected with *Mtb, Mtb*Δ*LprE* and *Mtb*Δ*LprE::LprE* strains for 3 h. Infected cells were harvested and checked for expression of **A**. EEA1 at 6 h, **B**. LAMP1, and Rab7 at 12 h post infection by western blotting. Uninfected cells were used as a control. Relative protein levels were plotted based on densitometry values. Expression of **C**. IL-12 and **D.** IL-22 was checked by qRT-PCR using gene specific primers at indicated time points. Expression of cytokines in uninfected THP-1 cells was normalized to 1 and fold change in infected cells was plotted. Experiments were performed in triplicates. Mean ± SD;*** for P<0.0001, ns, non-significant.

### LprE_Mtb_ arrest phago-lysosome maturation by down-regulating the expression of IL-12 and IL-22 cytokines

Upon mycobacterial infection cytokines strongly influence the outcome of infection. Cytokines individually or in combined fashion create a microenvironment that assists in the control of mycobacteria. Several cytokines have been shown to promote acidification of phagosomes to facilitate phago-lysosomal fusion and thus bacterial death. Other studies have shown that interleukin-12 (IL-12) and IL-22 cytokines play a critical role in phagosome maturation by promoting an acidification of the bacteria containing phagosomes [51,52]. IL-22 has been shown to enhance Rab7 expression in *Mtb* infected macrophages, thus enhancing phago-lysosomal fusion. Therefore, we investigated the expression of these two cytokines in infected THP-1 cells. As can be seen in **Figure 7C** transcript levels of IL-12 was elevated in a time dependent manner in macrophages infected with *Mtb*Δ*LprE* as compared to *Mtb* and *Mtb*Δ*LprE::LprE* strains. On the contrary, *Msm::LprE* infected cells showed reduced IL-12 expression as compared to vector control *Msm::pSMT3* infected cells (**Suppl Fig 6B**). The IL-22 level also increased by more than 2-fold in macrophages infected with the *Mtb*Δ*LprE* (**Fig 7D**), while *Msm::LprE* and *Msm*Δ*LprE::LprE* infected condition significantly reduced IL-22 transcript levels (**Suppl Fig 6C**). These results indicate that LprE_M_t_b_ probably blocks phagosomal maturation by down-regulating the expression of IL-12 and IL-22 cytokines.

## DISCUSSION

*Mtb* exhibits an extraordinary ability to survive inside the host cells. This is mainly attributed to the plethora of virulence factors produced by the bacterium. One of the most important classes of virulence factors produced by *Mtb* is lipoproteins. Of the 99 lipoproteins predicted in the mycobacterium genome, only a few of them have been functionally characterized. In this study, we have characterized one of the lipoproteins LprE, encoded by *Rv1252c* in H37Rv and *MT1291* in CDC1551 strains, and investigated its role in mycobacterial virulence. Analysis of genetic organization of *Mtb* genome showed that *Rv1252c* is a part of a previously uncharacterized Rv_dir301 operon. Sequence analysis revealed 100% homology between *lprE* gene from H37Rv and CDC1551 strains. LprE structural analysis showed the presence of alpha helices, random coils and N-terminal lipoprotein binding domain between 81-201 amino acids. Further domain analysis showed a perfect match between the binding domain sequences of LprE_Mtb_ and LptE lipoprotein from *E.coli*. The previous study by Sutcliffe *et.al*. also predicted LprE_Mtb_ as a lipoprotein on the basis of the presence of G+Lpp pattern match, Lipobox and type II signal peptide sequence [53]. These data provide sufficient evidences that LprE_Mtb_ could be a lipoprotein. However further biochemical analysis is required to confirm that LprE_Mtb_ is indeed a lipoprotein.

*Mtb* is considered as one of the most successful pathogens due to its ability to evolve with numerous survival strategies. However, non-pathogenic *Msm* is readily killed by macrophages. Here, we employed two strategies to understand the role of LprE in *Mtb* pathogenesis. First, we used a *Mtb* LprE deletion mutant (*Mtb*Δ*LprE*). Secondly, we used *Msm* as a surrogate model to express LprE_Mtb_ *(Msm::LprE).* To ensure that the observed phenotype is indeed due to ectopic expression of LprE in *Msm,* we deleted *MSMEG_5043*, a LprE_Mtb_ orthologue present in *Msm* genome, to generate *Msm*Δ*5043* strain. Infection assays with primary human macrophages and THP-1 cells showed that *lprE_Mtb_* expression increased the survival of *Msm*, while deletion of LprE significantly reduced the *Mtb* survival in macrophages suggesting that LprE_Mtb_ is involved in bacterial persistence in macrophages.

*Mtb* employs various immune evasion strategies to survive inside the macrophages. The antimicrobial peptide cathelicidin/LL-37 is known to play a significant role in innate response against *Mtb* infection [14,54]. Gutierrez *et al.* (2008) have shown that *Msm* infected macrophages differentially expressed a set of genes including cationic AMPs [55]. Our previous study has shown that *Msm* infection up-regulated the expression of cathelicidin in macrophages that led to the intracellular killing of bacteria. Moreover, we and others have shown that cathelicidin is able to kill *M. bovis*-BCG and *Mtb* H37Rv under *in-vitro and ex-vivo* growth conditions [13]. We observed that *Msm::LprE* and the complemented *Msm*Δ*5043::LprE* strains survived better in human macrophages by down-regulating the expression of cathelicidin, while deletion of *LprE* de-repressed the cathelicidin expression thereby resulting in increased *Mtb* killing. External supplementation of purified LL-37 peptide decreased *Msm::LprE* and *Msm*Δ*5043::LprE* bacterial counts confirming that the increased survival was due to LprE_Mtb_ mediated down-regulation of cathelicidin levels. Transcriptional down regulation of *CAMP* during bacterial infection was suggested as a novel mechanism to escape the immune responses [56,57,58]. LprE_Mtb_ was also found to inhibit the expression of cathelicidin signaling pathway intermediates such as CYP27B1 and VDR to subvert the Vitamin D3 mediated bacterial killing. Together, these findings argue that LprE_Mtb_ is responsible for the down-regulation of cathelicidin thus promoting bacterial survival in macrophages. Unlike LprE, LpqH_Mtb_ was shown to induce the expression of cathelicidin [16]. Infection assays with *Msm* strain co-expressing both LpqH_Mtb_ and LprE_Mtb_ (*Msm::LprE-LpqH*) showed intermediate survival and cathelicidin expression indicating that both LpqH_Mtb_ and LprE_Mtb_ are antagonistic regulators of cathelicidin expression.

One of the proposed mechanisms for cathelicidin induction is that bacterial exposure leads to activation of downstream p38 MAPK signaling pathway [16,54], which subsequently up regulates CYP27B1 hydroxylase, VDR and finally cathelicidin [16]. Indeed, our results demonstrated that *Mtb*Δ*LprE* infection caused up-regulation of p38 MAPK, CYP27B1, and VDR. On the other hand infection with LprE expressing *Mtb*, *Mtb*Δ*prE::LprE, Msm::LprE* and *Msm*Δ*5043::LprE* strains down-regulated p38 MAPK, CYP27B1, and VDR. These results suggest that LprE_Mtb_ mediate cathelicidin suppression via inhibition of p38-Cyp27B1-VDR signaling pathway. Previous reports have shown that few intracellular bacteria have the ability to dysfunction the vitamin D receptor, thus make host cells more susceptible to bacterial infections [59,60]. We observed that LprE_Mtb_ inhibits VDR and that treatment with an active form of Vitamin D3 further increased the killing of LprE_Mtb_ expressing mycobacterial strains indicating a possible mechanism of immunosuppression employed by the bacteria for intracellular survival.

Pro-inflammatory cytokine IL-1β is known for a protective immune action against mycobacteria [61]. IL-1β functions in synergy with LL-37 to augment the host immune responses against pathogenic bacteria [62]. Expression of LprE_Mtb_ decreased IL-1β levels, while deletion of LprE_Mtb_ was found to de-repress IL-1ß level in caspase-1 dependent mechanism. Thus, our data indicate that LprE_Mtb_ modulates IL-1β levels possibly in tandem with LL-37 downregulation to provide a favorable niche for bacterial survival. However, further mechanistic studies are required to establish the correlation between cathelicidin and IL-1β modulation by LprE.

Lipoproteins are known to trigger innate immune responses by binding to pattern recognition receptors, specifically TLR’s [63,64,65]. LprA, LpqH, and LprG are the three mycobacterial lipoproteins known to interact with TLR-2. Our current study has provided sufficient evidences that LprE_Mtb_ has a high binding affinity towards TLR-2 in comparison to TLR-1, 4 and 6. To further confirm the involvement of TLR-2, we checked bacterial survival and the expression of *CAMP*, *Cyp27B1* and *VDR* in TLR 1, 2, 4 and 6 silenced macrophages. We observed a significant increase in the expression of *CAMP*, Cyp27b1 and VDR that result in decreased survival of LprE expressing mycobacterial strains in TLR-2 silenced macrophages indicating that LprE_Mtb_ mediated inhibition of cathelicidin expression is a TLR-2 dependent phenomenon.

Many pathogenic bacteria, including *Mtb*, inhibit autophagy mechanism to facilitate its persistence in host cells [66]. Several bacterial virulence proteins were found to inhibit the autophagy. LpqH_Mtb_ activates autophagy, whereas EIS_Mtb_ (enhance intracellular protein) protein suppresses autophagy [67]. Our previous work also showed that a *Mtb* phosphoribosyltransferase suppressed autophagy expression to promote intracellular bacterial survival [32]. In the current study, we found that LprE_Mtb_ is also responsible for the suppression of autophagy as observed by the significant up-regulation of autophagic markers LC3, Atg-5, and Beclin-1 in cells infected with LprE_Mtb_ deletion mutant. These results indicate that LprE_Mtb_ suppress Vitamin D3 responsive cathelicidin, which may subsequently cause autophagic inhibition. Previous studies have shown that cathelicidin plays a role in the regulation of autophagy [68]. Altogether, inhibition of these antibacterial effector molecules by *Mtb* aids bacterial persistence inside macrophages.

One of the prominent mechanisms employed by *Mtb* to survive in macrophages is to block the phagosome maturation by inhibiting the recruitment of EEA1, Rab7, and LAMP-1, which are essential for the development of early and late endosomes and finally phagolysosome fusion. We have previously shown that exogenous administration of LL-37 increases the co-localization of *Msm* and *M. bovis*-BCG containing phagosomes with lysosomes [13]. Here, we found that infection with LprE_Mtb_ expressing strain decreased the expression of EEA1, LAMP1, and Rab7 in THP-1 cells, while *Mtb*Δ*LprE* infection induced the expression of these proteins indicating that LprE_Mtb_ supports bacterial survival by preventing phago-lysosome fusion. *L. monocytogenes* lipoproteins have also been shown to play an important role in the phagosomal escape of bacteria [69].

Stimulation of macrophages with cytokines restricts the growth of microorganisms within endocytic compartments [70]. It is shown that treatment of human macrophages with IL-12 restrict the growth of *M. bovis*-BCG by facilitating the phago-lysososome fusion [51]. Similarly, treatment with IL-22 was shown to increase the expression of endocytic marker proteins and increased bacterial killing by inducing phago-lysosomal fusion [52]. In our study, we observed decreased levels of IL-12 and IL-22 in cells infected with LprE_Mtb_ expressing mycobacteria, while expression of these cytokines was found to be increased in *Mtb*Δ*LprE* infected cells. These results indicate that LprE_Mtb_ mediates phagosomal arrest by down-regulating the expression of IL-12 and IL-22 cytokines. Moreover, lower levels of cathelicidin and IL-1β also explain the inhibition of phagosome maturation and fusion with lysosomes.

Collectively, our findings identified a new *Mtb* lipoprotein that could be used by pathogenic mycobacteria to alter host immune responses thus presenting attractive targets for new drug therapies. **Figure 8** shows a schematic representation of modulation of different host immune responses by LprE_Mtb_ leading to increased bacillary survival in macrophages.

**Fig 8:**
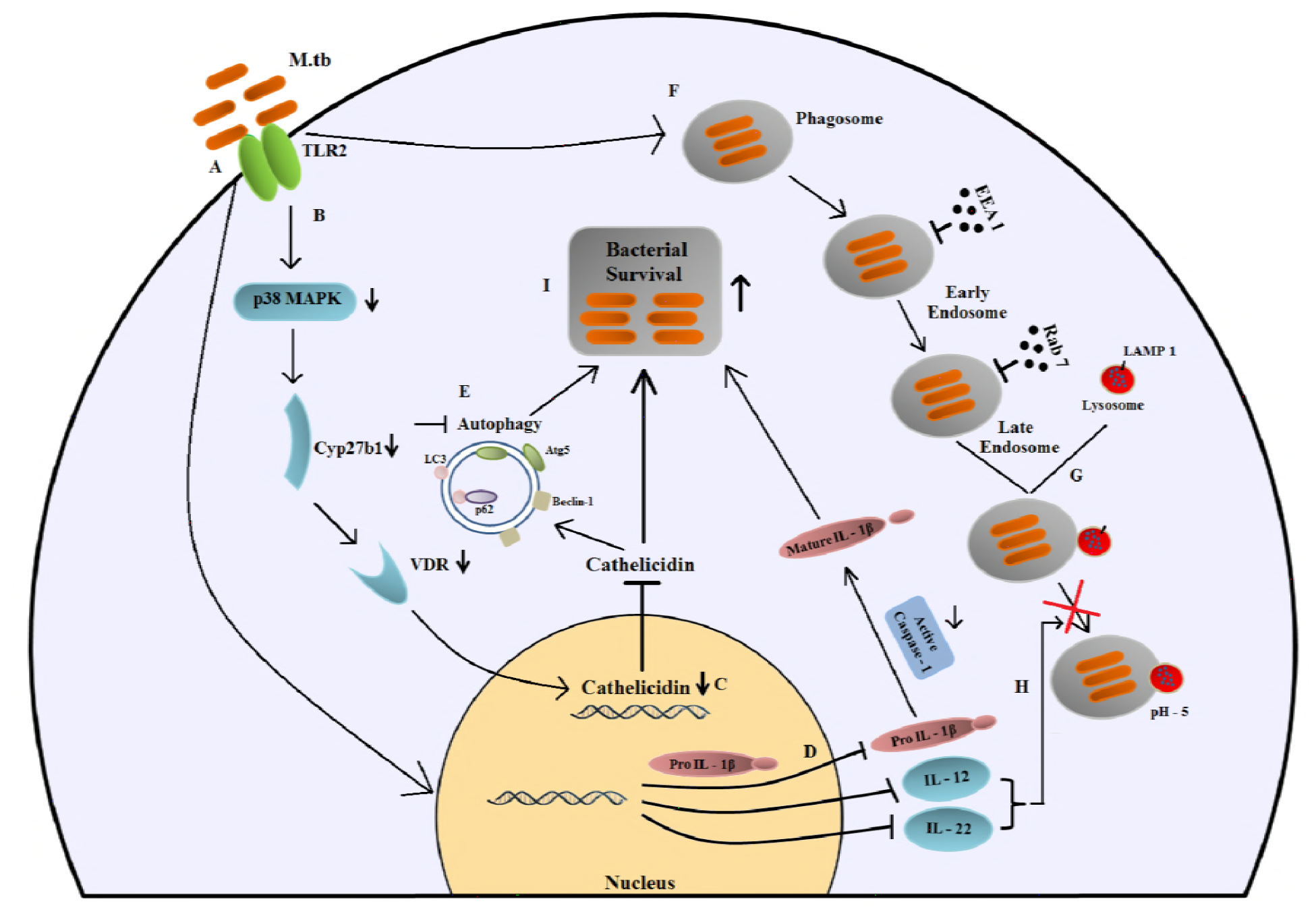
Schematic representation of LprE_Mtb_ mediated immune modulation leading to increased bacterial survival in macrophages. **A**. LprE mediates the immune response through interaction with TLR-2 of macrophages, **B**. which then down-regulates expression of p38-Cyp27B1-VDR signalling pathway and **C**. *CAMP* expression leading to increased bacterial survival. **D**. **LprE**_Mtb_ mediated down-regulation of antimicrobial action of caspase-1 dependent IL-β. **E**. Inhibition of autophagic markers leading to increased bacterial survival. **F**. **LprE**_Mtb_ inhibits the recruitment of EEA1 and RAb7 to early stages of endosome formation and G. LAMP1 recruitment occurs at later stages thus arresting the phago-lysosomal fusion and this arrest is mediated by decreased **H**. IL-12 and IL-22 cytokine levels. **I**. All these facets of host immune modulation by **LprE**_Mtb_ lead to significant increase in intracellular bacterial survival.

## MATERIALS AND METHODS

### Ethics and Biosafety Statement

All experiments were approved by the Institutional Biosafety committee of KIIT University (vide DBT memorandum No-BT/BS/17/493/2012-PID). All the bacterial mutants were handled in adherence to experimental guidelines and procedures approved by the Institutional Biosafety Committee (IBSC) of School of Biotechnology, KIIT University (KIIT/3-12).

### *In-silico* analysis

Bacterial gene sequences were obtained from Tuberculist (http://tuberculist.epfl.ch/) and Uniprot (http://www.uniprot.org/) databases. Protein secondary structure was predicted using SOPMA secondary structure based method (https://npsa-prabi.ibcp.fr/cgi-bin/npsa_automat.pl?page=/NPSA/npsa_sopma.html). For tertiary structure prediction, LprE_Mtb_ (Rv1252) sequence was BLAST against Protein Data Bank (PDB) to obtain a suitable template. Structure showing highest homology with human mitochondrial RNA polymerase (PDB ID: 3SPA) was chosen as a suitable template. Using this template, tertiary structure of LprE_Mtb_ was depicted using Modeller9.12 [71]. The modelled LprE_Mtb_ structure was energy minimized and validated using ModRefiner (http://zhanglab.ccmb.med.umich.edu/ModRefiner/) and RAMPAGE (http://mordred.bioc.cam.ac.uk/~rapper/rampage.php) web based servers. LprE_Mtb_ domain analysis was done using Pfam (http://pfam.xfam.org/) database and visualized using PyMOL software v1.7.4 (http://www.pymol.org). Human toll like receptor-1,2, 4 and 6 structures were obtained from Protein Data bank (http://www.rcsb.org/pdb/home/home.do). Molecular docking of the structures was done using PatchDock server (http://www.bioinfo3d.cs.tau.ac.il/PatchDock/) and visualized using PyMOL. The Atomic surface energy was determined to analyze the protein-protein binding intensity.

### Bacterial strains, cell lines, and reagents

*M. tuberculosis* CDC1551 *(NR-13649)* and *M. tuberculosis* CDC1551 LprE mutant (NR-17996) were obtained from BEI Resources, USA. All *Mtb* strains were grown in Middlebrook 7H9 broth or on Middlebrook 7H10 plates (Difco, Sparks, Maryland, USA) as described previously [32–34]. *Escherichia coli* XL-10 Gold (Stratagene, San Diego, California, USA) was grown in Luria-Bertani (LB) broth supplemented with 20 μg/ml tetracycline. pSMT3 vector was a kind gift from Dr. Rakesh Sharma (IGIB, Delhi). THP-1 cells were maintained in RPMI Glutamax (Gibco, Waltham, MA, USA) medium supplemented with 10% fetal bovine serum (FBS) (Gibco) and penicillin-streptomycin solution (Gibco) at 37 ^0^C and 5% CO2.Vitamin D3 and tetracyline were purchased from Sigma Aldrich (USA). TRIzol was purchased from Invitrogen (Carlsbad, California, USA). Complementary DNA (cDNA) synthesis kit was procured from Fermentas (USA). Trypsin-EDTA, SDS, Tris-HCl, and NaCl were obtained from Sigma-Aldrich (St. Louis, Missouri, USA). siRNA oligos were designed and manufactured by GeneX India Bioscience Pvt. Ltd (Chennai, India). All the cloning reagents and restriction enzymes were obtained from NEB (Ipswich, MA, USA).

### Construction of *M. tuberculosis* CDC1551 LprE strain

For the construction of complemented strain, *LprE* was amplified from Mtb CDC1551 genomic DNA using gene specific primers (**Table 1**) and was cloned into *NdeI-Hind III* sites of the pVV16 vector [72] and pNiT vector [72]. The pVV16-LprE and pNit-LprE constructs were transformed into *MtbΔLprE* mutant to generate *MtbΔLprE::LprE* complemented strain.

**Table 1:**
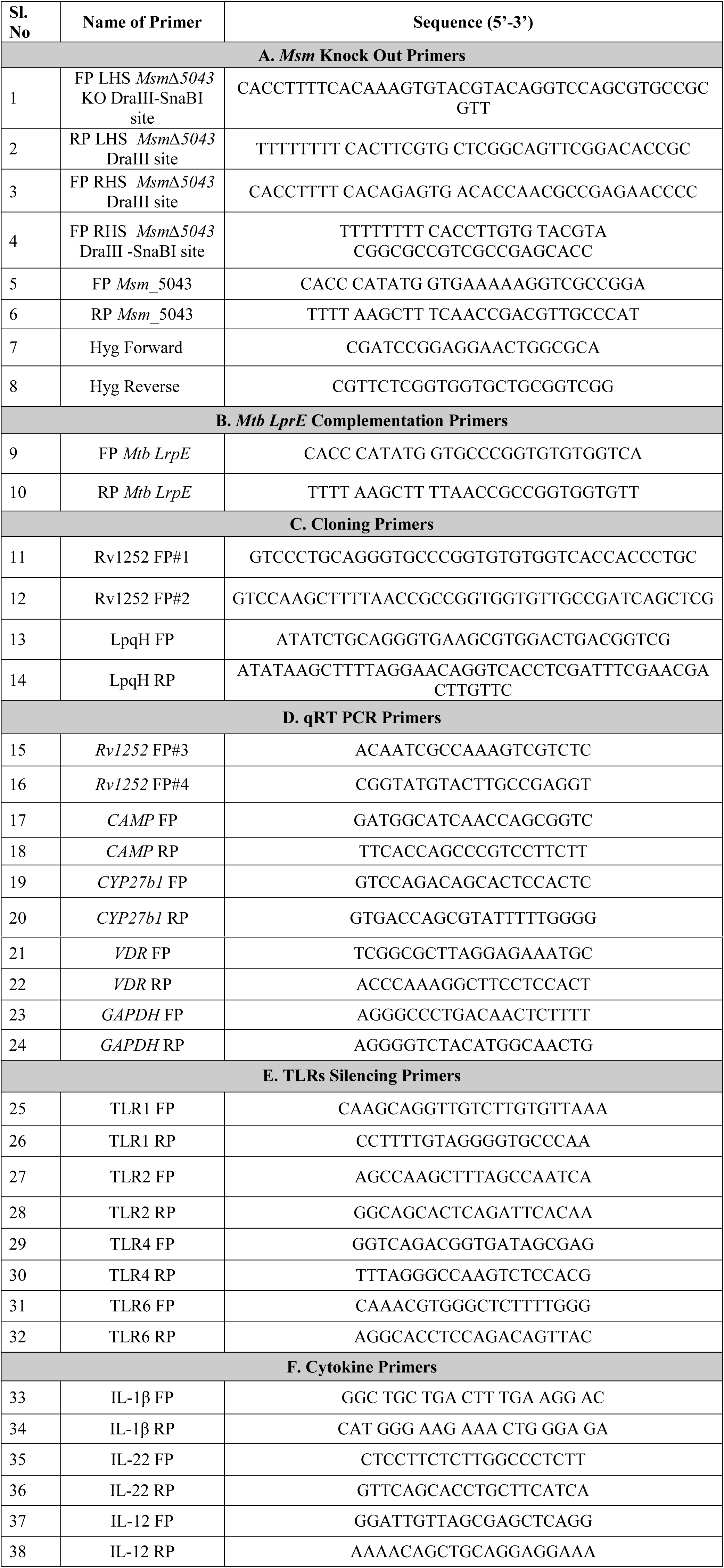
**List of primers used in this study.** FP: Forward primer, RP: Reverse primer, LHS: Left hand site, RHS: Right hand site.

### Construction of recombinant *M.smegmatis* strain expressing *LprE_Mtb_ and LpqHMt_b_*

*LprE_Mtb_* and *LpqH_Mtb_* were PCR amplified using gene specific primers (**Table 1**) using genomic DNA as a template. The PCR amplified products were purified from the gel, double digested with *Pst*I and *Hind*III and cloned into pSMT3 shuttle vector separately. The recombinant constructs were transformed into competent *E. coli* XL-10 gold. LB agar plates supplemented with 20 μg/ml tetracycline and 50 μg/ml hygromycin was used to select the positive colonies and were further confirmed by colony PCR and sequencing using gene specific primers. Finally, the recombinant constructs were transformed into electro competent *Msm*. The positive colonies were selected on 7H10 medium containing 50 μg/ml hygromycin. The positive transformants were confirmed by colony PCR and sequencing using gene specific primers (**Table 1**).Generation of recombinant *Msm::LprE-LpqH* strain was confirmed by sequencing using gene specific primers.

### Construction of *M. smegmatisΔ5043* mutant

The allelic exchange substrate (AES) for the generation of *Msm 5043* mutant (*Msm*Δ*5043*) was produced as described previously [72]. Briefly, approximately 800 kb regions upstream and downstream of *MSMEG_5043* loci were PCR amplified using specific primers (**Table 1**). The amplicons were digested with DraIII-HF (NEB) and ligated with two compatible fragments (OriE+cosλ and Hyg^res^) from pYUB1474 [73] and pVV16 [72], respectively to generate allelic exchange substrate (AES). *MSMEG_5043-AES* was linearized with *SnaBI* to elute upstream-Hyg-downstream flank. AES was electroporated into electro-competent *Msm* cells. Hygromycin-resistant colonies were screened by PCR to confirm deletion of *MSMEG_5043* at its genomic locus (*Msm*Δ*5043*). Then *LprE-pSMT3* construct was electroporated into *Msm*Δ*5043* mutant to create a complemented *Msm*Δ*5043::LprE* strain.

### Growth kinetics

*Msm::pSMT3* vector control, recombinant *Msm* expressing LprE_Mtb_ (*Msm::LprE*), *Msm*Δ*5043* and *Msm*Δ*5043::LprE* strains were grown in 7H9 broth at 120 r.p.m at 37^0^C with. The growth kinetics was assayed by measuring the O.D at 600 nm (OD_600_) at regular intervals till death phase was observed. Similarly growth kinetics of *Mtb, MtbΔLprE* and *Mtb*Δ*LprE::LprE* was determined by measuring OD_600_ at regular intervals till death phase was observed.

### Intracellular survival assay

*Mtb*, *Mtb*Δ*LprE*, and *Mtb*Δ*LprE::LprE, Msm::pSMT3, Msm::LprE, Msm*Δ*5043* and *Msm*Δ*5043::LprE* strains were grown to mid-exponential phase. The bacterial cultures were pelleted at 5000 g for 5 min, washed twice with 1X phosphate buffer saline (PBS) and resuspended in RPMI medium to adjust the OD_600_ to 0.1. THP-1 monocytes (2x10^5^cells/well) were differentiated into macrophages by addition of 20 nM phorbol 12-myristate 13-acetate (PMA) and incubated for 24 h. After which the cells are washed with RPMI medium to remove PMA, and then the cells are allowed to differentiate for 3 days at 37 ^0^C and 5% CO_2_. Cells were infected as described previously at a multiplicity of infection (MOI) 10 [74, 32]. The intracellular bacterial survival was determined by lysing the cells with chilled 0.5% Triton-X 100 (Sigma) at different time points and plating the serially diluted samples on 7H10 medium. Bacterial colonies were enumerated after 3 and 21 days for *Msm* and *Mtb*, respectively.

### Flow cytometry analysis

THP-1 cells (2x10^5^ cells/well) were seeded on a 24-well plate and treated with PMA as described above. Then cells were infected with *Msm::pSMT3*, *Msm::LprE*, *Msm*Δ*5043*, and *Msm*Δ*5043::LprE* at an MOI 10 as described above. After infection, cells were lysed by treating with 0.5% Triton X-100 for 2 min to release the intracellular bacteria. The cell suspension was centrifuged at 1000 g for 10 min to pellet down the lysed macrophage cells. Supernatants that contained released bacteria were again centrifuged at 12000 g for 10 min. The pellet was washed thrice with 1X PBS, bacterial cells were stained with 30 μM propidium iodide (PI) (Thermo Fisher, P3566) for 5 min in dark and the live cells were analyzed using BD FACSCantoII flow cytometer. Threshold (n=1000) was set on Side Scatter (SSC), and PMT voltages were set using an unstained sample of diluted bacteria grown to log phage. The bacterial population was positioned so that the entire population is on the scale on an FSC vs SSC plot. A total of 10,000 events are acquired and a gate was set according to the stained bacterial sample. Uninfected THP-1 cells stained with PI were also used to rule out cell contamination while analysis.

### Phagocytosis assay

THP-1 cells (2x10^5^ cells/well) were infected with *Msm::pSMT3*, *Msm::LprE*, *Msm*Δ*5043*, and *Msm*Δ*5043::LprE* at an MOI 10. 1 and 2 h post infection cells were washed rigorously with 1X PBS to remove un-internalized and membrane adhering bacteria. The cells were lysed with 0.5% Triton X-100 and released intracellular bacteria were plated onto 7H10 plates to count live bacteria. Bacterial preparations were plated before infection to ensure that an equal number of bacteria were used for infection assays.

### Isolation of human peripheral blood mononuclear cells (PBMCs)

Blood was collected from healthy individuals, diluted (1:1 in PBS), layered onto an equal volume of Ficoll-Paque plus (GE Healthcare, 17-1440-02) and centrifuged at 100 g for 30 mins. The whitish buffy coat was washed twice with 1x PBS (100Xg, 10 mins). The cell pellet was resuspended in RPMI medium supplemented with 10% FBS. 1x10^5^ cells were seeded onto 24-well for 24 h at 37 °C. Fresh RPMI media containing 5 ng/ml GM-CSF (R&D Systems, McKinley, MN, USA) was added per well and maintained at 37 °C for 72 hours. After differentiation, the cells were infected with mycobacterial strains as described above.

### LL-37 treatment

LL-37 peptide (LLGDFFRKSKEKIGKEFKRIVQRIKDFLRNLVPRTES) was synthesized (GeneXbio, Delhi, India) at a purity of >95%. The peptides were dissolved in 0.01% acetic acid and stored at -20°C until further use. THP-1 cells were treated with LL-37 (50μg/ml) 2 h before (pre-treated) and after (post-treated) infection as described previously [11].

### TLR silencing

siRNA duplex against TLR-1, 2,4 and 6 were designed and synthesized (GeneX, New Delhi, India) at 10 nmol concentration. 20 nM siRNAs were electroporated into 10^6^ THP-1 cells. Scrambled siRNA was used as a negative control. The electroporated cells were treated with 20 nM PMA and seeded onto 6 well plates for infection assays. Silencing efficiency was determined using gene specific primers.

### Western blotting

All the antibodies except LC3 and CYP27B1 were purchased from Cell Signaling. The expression of phospho-p38 (Cat No-4511), total p38 (Cat No-9212), β actin (Cat No-4970),CYP27B1 (Thermo Scientific, Cat No-CA5 26065), VDR (Cat No-12550), Rab7 (Cat No-2094), EEA1(Cat No-3288), LAMP1 (Cat No-9091), LC3 (Sigma, L7543), SQSTM1/p-62 (Cat No-5114), Atg-5 (Cat No-8540) and Beclin-1 (Cat No-3495) was determined in THP-1 cells infected with *Mtb*, *MtbALprE*, *MtbALprE::LprE*, *Msm::pSMT3*, *Msm:;LprE*, *Msm*Δ*5043* and *Msm*Δ*5043::LprE* strain as described above. Cells were harvested at the indicated time points by adding 80-100 μl lysis buffer (1M Tris, 2M Nacl, 0.1M EDTA, 100 mM DTT, 1% Triton X100, NasVO4'2H2O, 10% glycerol and PMSF) and stored in -80°C overnight. The lysates were thawed on ice, briefly vortexed for 30 secs for three times and centrifuged at 13,000 g. for 30 mins. The isolated proteins were estimated by Bradford assay, run on SDS-PAGE, electrophoresed and transferred onto PVDF membrane (GE healthcare, Chicago, Illinois, USA) for 90 mins at 50 mA. Then the blots were blocked with either 5% BSA or 5% skimmed milk for 2 h at room temperature. The membranes were then incubated with primary antibodies (1:1000) overnight at 4 °C. The membranes were then washed thrice with 1X PBS containing 1% Tween 20 (PBST) for 5 min. The washed membrane was then incubated with goat anti-rabbit IgG HRP secondary antibody (GeneI merck, 6.2114E14)) for 2 h at room temperature. The membranes were then washed with 1X PBST for three times 5 min each and developed on X-ray film using chemiluminescent solvents. The band densities were quantified by computer scanning of the films and normalizing the values for analysis. Image J2 (NIH, USA) was used for computer analysis of pixel intensity of bands on films. Relative band densities relative to respective loading control were determined.

### Quantitative real time PCR

To check the expression of *LprE_Mtb_* as a function of growth, *Msm::pSMT3* and *Msm::LprE* strains were grown in 7H9 medium. Bacteria were harvested at different time points. Total RNA was isolated by using TRIzol reagent and cDNA was synthesized using Verso cDNA synthesis kit as per the manufacturer’s instructions (Thermo Scientific). qRT-PCR amplification was carried out using gene specific primers and the synthesized cDNA as a template. All reactions were performed in a total reaction volume of 10 μl using SYBR^®^ Green PCR mastermix (KAPA Biosystems), and carried out in Insta Q96 (Himedia, India) with initial denaturation at 95°C for 10 min, final denaturation at 95°C for 30 secs, annealing at 55°C for 30 sec and extension at 72°C for 30 sec to generate 200 bp amplicons. Similarly, the expression of *CAMP*, *Cyp27b1*, and *VDR*, *IL-12*, *IL-22 and IL-1β* were measured by isolating RNA from infected THP-1 cells at mentioned time points post infection using gene specific primers (**Table 1**). The reaction conditions were as follows; initial denaturation at 95 °C for 10 min, final denaturation at 95°C for 30 secs, annealing at 55°C for 30 secs, and extension at 72°C for 30 sec. The mRNA levels were normalized to the transcript levels of *sigA* and GAPDH and the relative fold changes were calculated. All quantitative PCR experiments were performed for three replicates and the data represented for each sample were relative to the expression level of housekeeping gene calculated using the 2^-ΔΔCT^ method.

### Statistical analysis

Statistical analysis was performed with GraphPad Prism v 5.0 (GraphPad Software, La Jolla, California,USA, http://www.graphpad.com). Results are expressed as mean ± SD, unless otherwise mentioned. Significance was referred as *** for P < 0.0001, ** for P < 0.001, * for P < 0.05.

## ACKNOWLEDGEMENTS

The Senior Research Fellowship (80/902/14-ECDI) awarded to Avinash Padhi from Indian Council of Medical Research (ICMR), Govt. of India is highly acknowledged. We are thankful to all members of Avinash Sonawane, Vinay Nandicoori and Pawan Gupta laboratory for valuable discussions.

## SUPPLEMENTARY FIGURES

**Suppl Fig 1:**
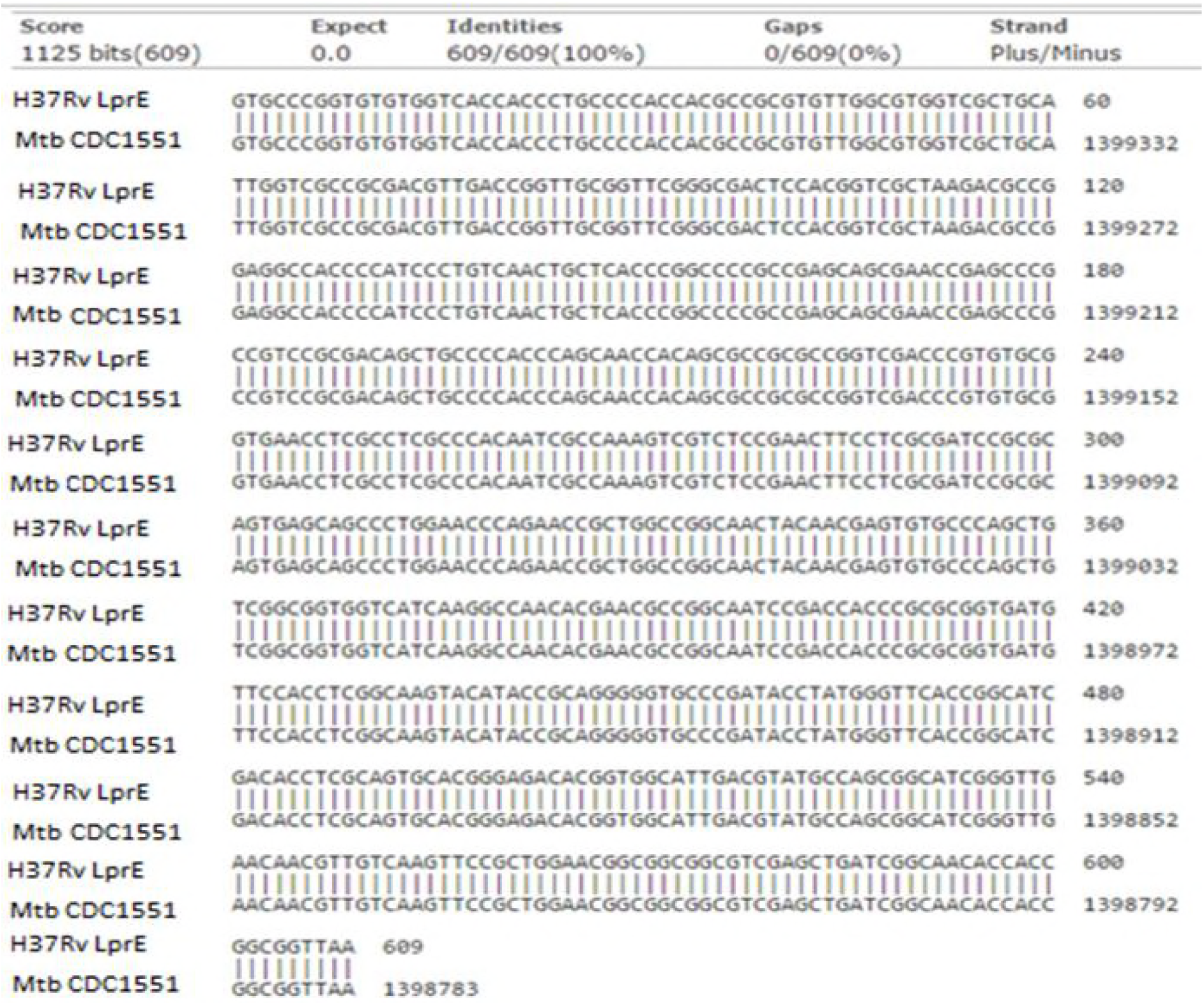
Multiple sequence alignment of *Mtb* H37Rv and CDC1551 LprE.

**Suppl Fig 2:**
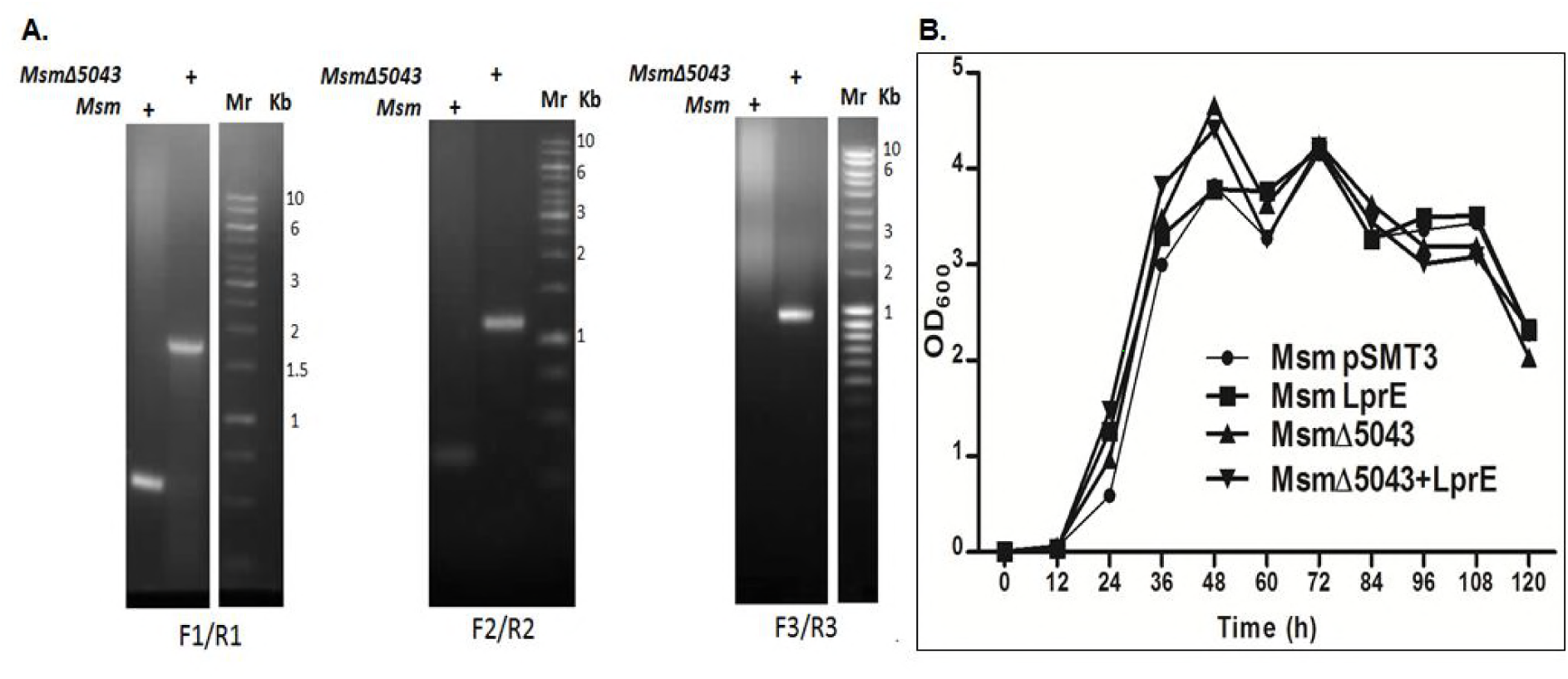
**A**. Disruption of *LprE_Mtb_* at a native locus was confirmed by performing PCRs using gene specific primers (F1/R1), beyond flank forward (F2) with Hygromycin reverse (R2) and Hygromycin forward (F3) with beyong flank reverse (R3). Genomic DNA from *Msm* (Lane 1) and *Msm*Δ*5043* (Lane 2) were used as a template. Mr represents 1 kb gene ruler ladder. **B**. *In vitro* growth curve of *Msm::pSMT3*, *Msm::LprE*, *Msm*Δ*5043* and *Msm*Δ*5043::LprE* was determined by growing bacteria in 7H9 medium and measuring optical density at A_600_ nm.

**Suppl Fig 3:**
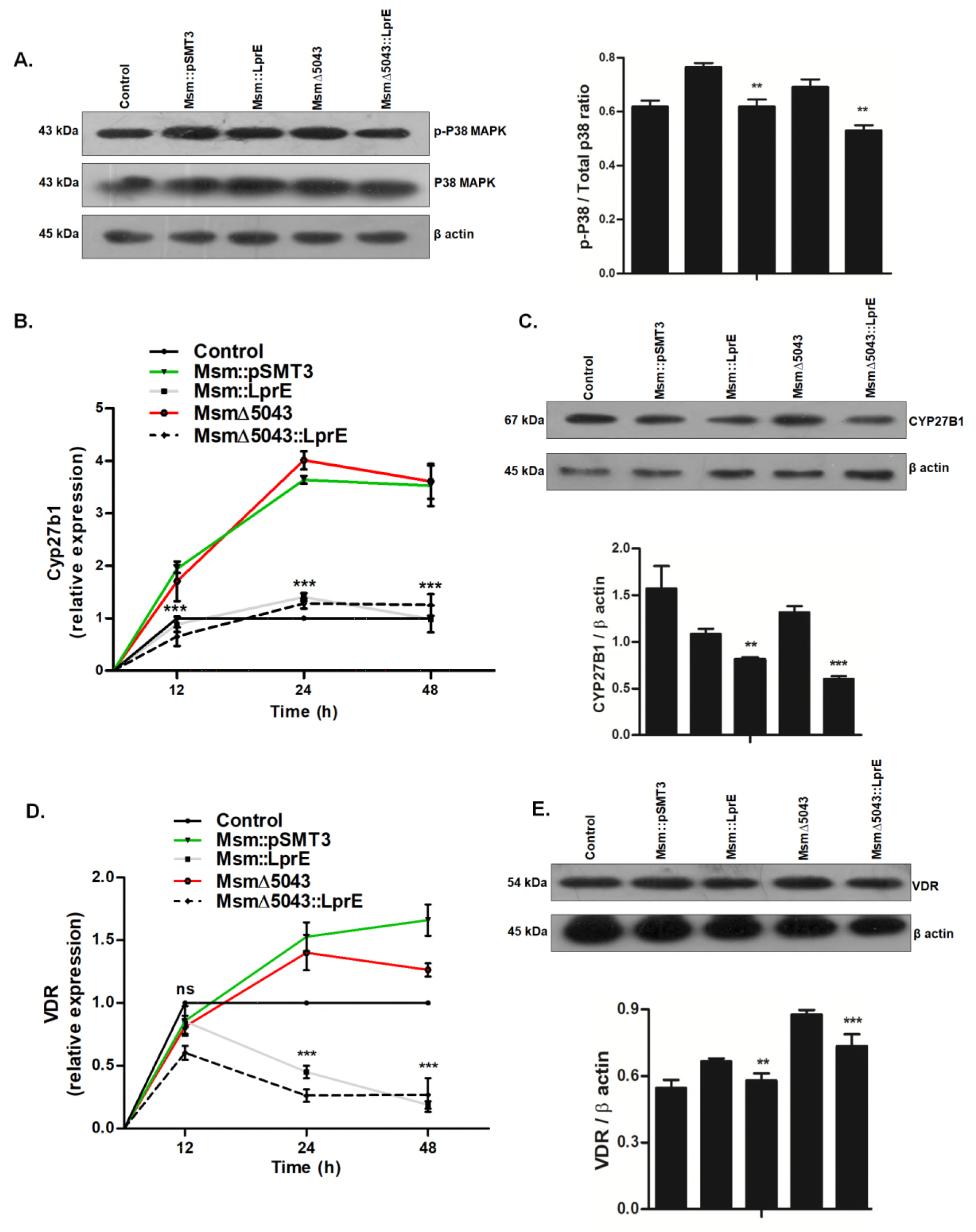
LprE_Mtb_ overexpression in *Msm* mediates cathelicidin downregulation through p38–Cyp27B1–VDR pathway. THP-1 cells were infected with *Msm::pSMT3*, *Msm::LprE*, *Msm*Δ*5043* and *Msm*Δ*5043::LprE* strains. **A**. Western blot analysis of phospho-p38 MAPK expression was performed using anti-p38 antibody 6 h post infection. Expression of *CYP27B1* in *Msm::pSMT3* infected THP-1 cells was determined 24 h post infection at **B**. transcriptional level by qRT-PCR and **C**. translational level by western blotting. Expression of VDR at **D**. transcriptional and **E**. translational level was determined by qRT-PCR and Western blotting, respectively in infected THP-1 cells. Experiments were performed in triplicates; Mean ± SD; *** for P < 0.0001, ** for P < 0.001; ns, non-significant.

**Suppl Fig 4:**
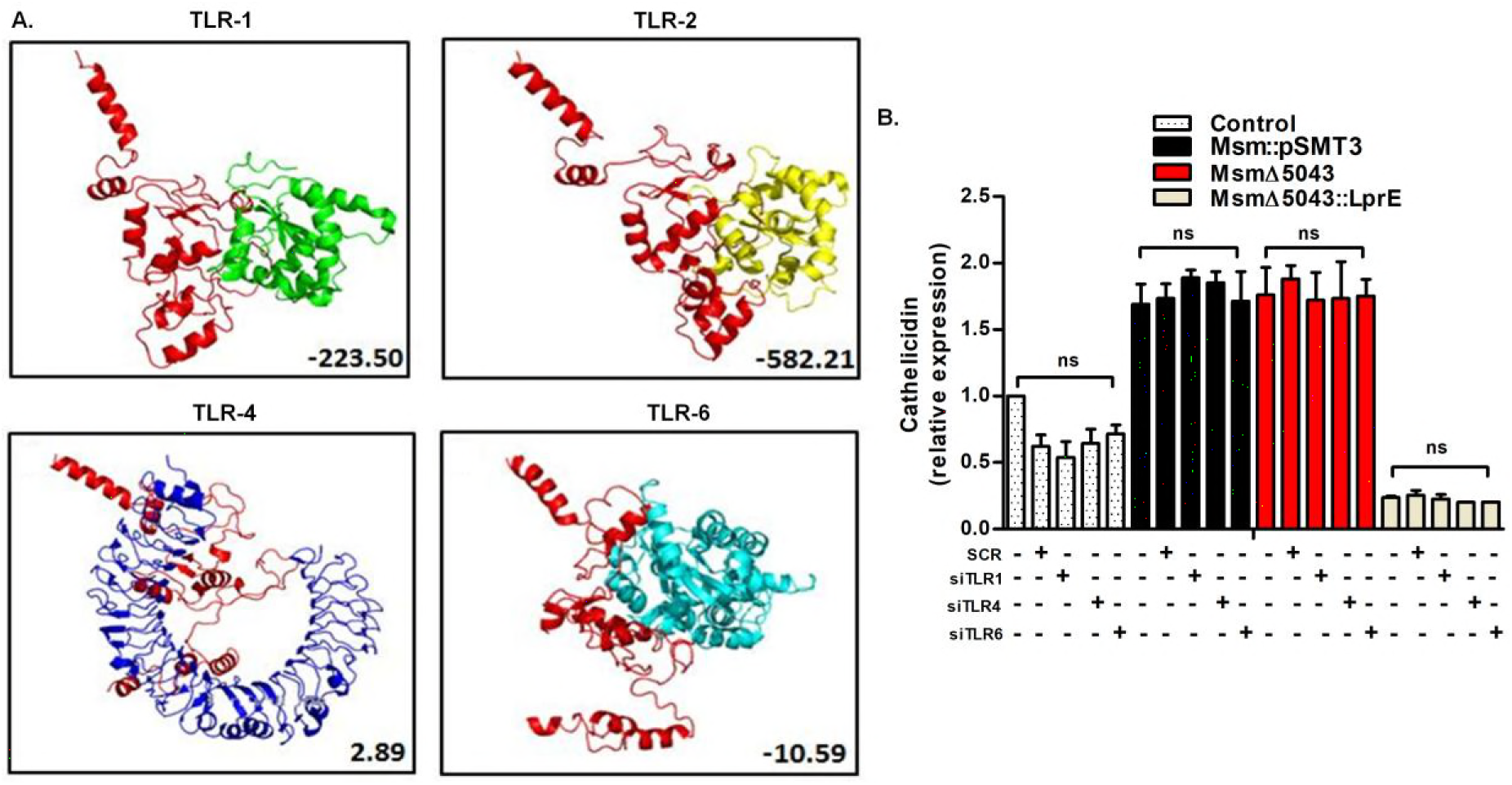
**A**. *In-silico* docking of energy minimized structure of LprE_Mtb_ with human TLR-1, TLR-2, TLR-4, and TLR-6. Numbers mentioned below in respective boxes indicate atomic contact energy (ACE) score obtained from PatchDock analysis **B**. Cathelicidin (*CAMP*) expression was checked by qRT-PCR in untreated, scrambled and si RNA against TLR-1/4/6 treated cells. Experiments were performed in triplicates; ns, non-significant.

**Suppl Fig 5:**
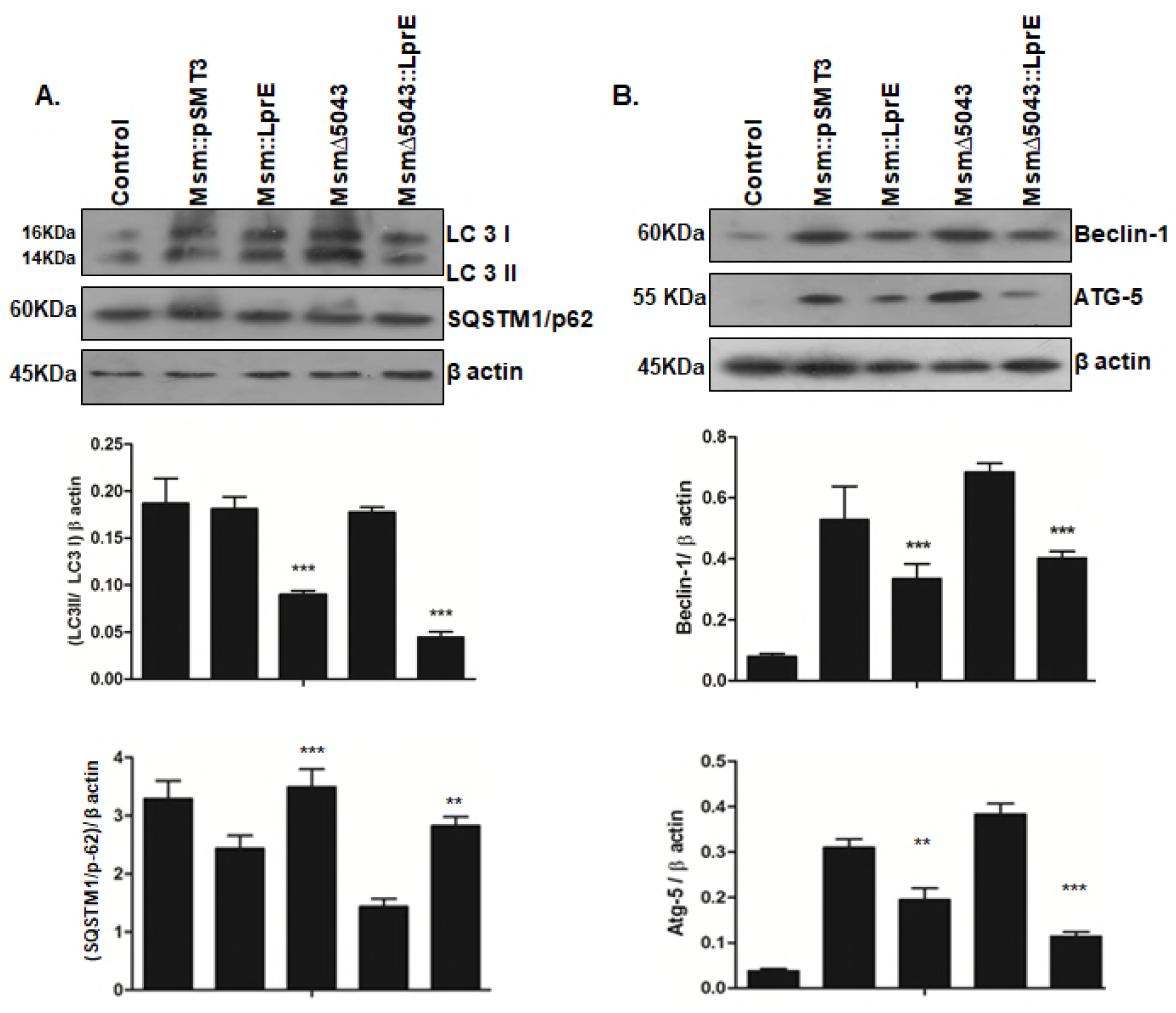
THP-1 cells were infected *Msm::pSMT3*, *Msm::LprE*, *Msm*Δ*5043* and *Msm*Δ*5043::LprE* strains. Western blotting analysis of **A.** LC3, SQSTM1/p62 at 6 h and **B.** Atg-5, Beclin-1 was done at 24 h post infection. Quantitative densitometry of expression was done using Image J software under baseline conditions. The expression values were normalized with β actin in western blotting. The blots are representative of three independent experiments; Mean ± SD; *** for P < 0.0001, ** for P < 0.001; ns, non-significant.

**Suppl Fig 6:**
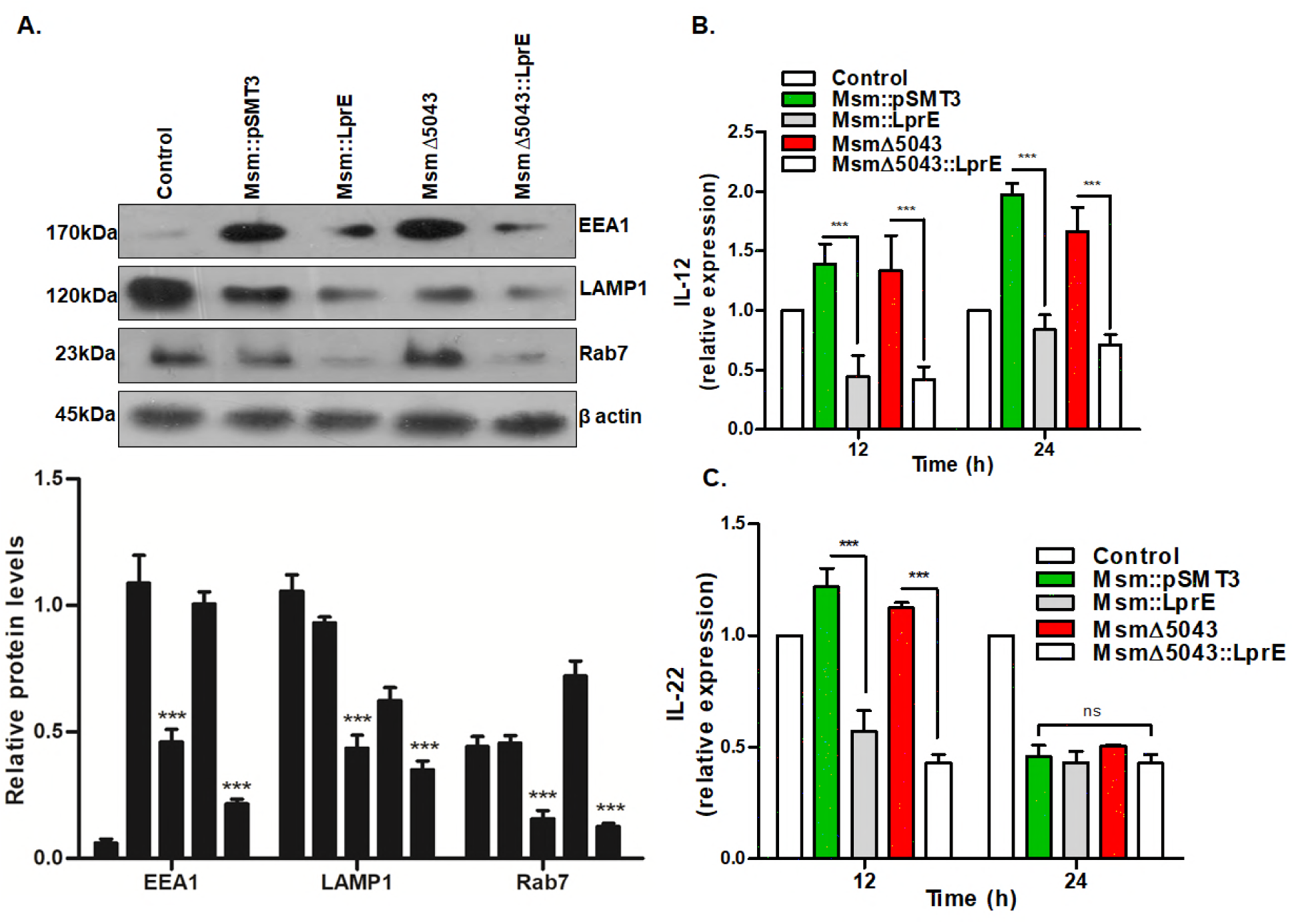
THP-1 cells were infected with *Msm::pSMT3*, *Msm::LprE*, *Msm*Δ*5043*, and *Msm*Δ*5043::LprE* strains. Infected cells were harvested to isolate proteins and total RNA. **A.** Western blotting was done to check for expression of EEA1, Rab7, and LAMP1. Un-infected cells were taken as a control, and densitometry of protein levels relative to β actin was plotted (below). RNA isolated at indicated time points was converted to cDNA and the expression of **B.** IL-12 and **C.** IL-22 was checked by qRT-PCR. The expression levels were normalized against GAPDH as the house keeping gene. Experiments were performed in triplicates. Mean ± SD; *** for P<0.0001, ns, non-significant.

